# Macro-Chemical Biology: Engineering Biomimetic Trogocytosis with Farnesylated Chemically Self-Assembled Nanorings

**DOI:** 10.1101/2022.03.01.482559

**Authors:** Yiao Wang, Lakmal Rozumalski, Caitlin Lichtenfels, Jacob R. Petersberg, Ozgun Kilic, Mark D. Distefano, Carston R. Wagner

**Affiliations:** Department of Chemistry, University of Minnesota, Minneapolis, Minnesota 55455, USA; Department of Medicinal Chemistry, University of Minnesota, Minneapolis, Minnesota 55455, USA

## Abstract

With the recent success of cell-based therapies, there has been a rapidly emerging interest in the engineering of cell-cell interactions and communications. Inspired by the natural intercellular material transfer process of trans-endocytosis or trogocytosis, we proposed that targeted farnesylated chemically self-assembled nanorings (farnesyl-CSANs) could serve as a biomimetic trogocytosis vehicle for engineering directional cargo transfer between cells; thus, allowing cell-cell interactions to be monitored, as well as facilitating communication between the cells by delivery of bioactive species. The membranes of sender cells were stably modified by hydrophobic insertion with the targeted farnesyl-CSANs and to be efficiently transferred to receiver cells expressing the appropriate receptor by endocytosis. CSAN-assisted cell-cell cargo transfer (C4T) was demonstrated to be receptor-specific and dependent on direct cell-cell interactions, the rate of receptor internalization and the amount of receptor expression. In addition, C4T was shown to facilitate cell-to-cell delivery of an apoptosis inducing drug, as wells as antisense oligonucleotides (ASO). Taken together, the C4T approach is a potentially versatile biomimetic trogocytosis platform that can be used to monitor cell-cell interactions, as well as the engineering of cell-cell communications, such as cell-based drug delivery.

## Introduction

Multicellular life and disease are dependent on the engagement of cells with each other, either for the development, maintenance and regeneration of tissues or identifying and removing diseased tissues^1–6^. Cell-cell interactions between glial cells and neural cells are key to central nervous system (CNS) functions^7,8^, while the contacts between T-cells and peripheral tissues are essential for defending against viral infections^9,10^. Cell-based therapeutics have rapidly emerged and expanded as invaluable tools in translational medicine with a significant impact on several diverse fields, including tissue engineering, regenerative medicine, and immunotherapy^11–13^. In each of these cases, cell-cell interactions, whether natural or genetically engineered, are key to understanding the biological processes wholistically, as well as assessing clinical success. In addition, the potential to augment and modulate the effects of cells on other cells is of key interest to engineering synthetic biological processes. Consequently, recent approaches have begun to be developed for the monitoring and engineering of cell-cell interactions. Cell-cell interactions have been monitored by genetically engineering cells to take advantage of surface modifying non-discriminating chemical or conjugation reactions or non-genetically by microdissection methods^14,15^. Genetic cargo transfer based approaches have emerged that rely on the binding of engineered receptors on receiver cells to engineered fluorescent or chemically modified proteins fused to membrane spanning domains on sender cells^16,17^. In this last approach, cargo transfer has also been used to deliver proteins and nucleic acids to the receiver cells from the sender cells^16^. Each of these approaches has proved versatile. Nevertheless, in each case genetic engineering of either the sender cell or receiver cell or both is required, which can be time consuming and, in some cases, difficult, since not all cells are amenable to genetic modification^18^. Consequently, alternative methodologies that allow for the non-genetic modification of normal cells and the evaluation of their interactions with cells expressing a variety of natural or engineered receptors would be of value. In addition, the potential for modulation of the receiver cell biology through a non-diffusible modulating ligand from another cell, would expand and complement diffusible cell-cell communications approaches; thus, facilitating interrogation and control of cell-cell interactions both in healthy and diseased tissues.

Previously, we reported the farnesylated CSANs as a universal system to modify mammalian cell surface for reversible cell-cell interactions^19^. The CSANs are oligomerized into predominantly octameric nanorings through the self-assembly of DHFR^2^ fusion proteins by a chemical dimerizer, bisMTX^20–22^ (Fig. 1a). The targeting fragments incorporated to the N-terminus of the DHFR^2^ proteins enable the CSANs to engage specific cellular receptors, while the C-terminal “CVIA” sequence of the proteins can be recognized and rapidly farnesylated by protein farnesyaltransferase^23–30^. The consequent farnesylated DHFR^2^ proteins can be easily self-assembled into farnesylated CSANs (farnesyl-CSANs) by incubation with bisMTX and used to efficiently modify mammalian cell membranes through the hydrophobic interactions between the isoprenoids groups of the nanorings and the membrane phospholipids^19^ (Fig. 1b). Farnesyl-CSANs were shown to be stably bind to cell surfaces for days (T_1/2_ > 3 days) and direct modified cells to the desired target cells^19^. In this study, we report that the farnesyl-CSANs on sender cell membranes are able to undergo efficient CSAN-assisted cell-cell cargo transfer (C4T). C4T was found to be dependent on the internalization rate of the surface receptor on the receiver cell and the amount of the targeted receptor expressed on the receiver cell. In addition, by varying the concentration of the farnesyl-CSANs on the sender cells, induced interactions between the sender and receiver cells could be kept at a minimum, thus reducing the effect of the modification on natural or engineered interactions between the sender and receiver cells. Moreover, by incorporating the azide-functionalized farnesyl analogs as bioconjugation handles^31^, both the apoptosis inducing drug, MMAE, and antisense oligonucleotide targeting expression of the translation initiation factor, eIF4E, were able to undergo C4T transfer (Fig. 1c). Thus, the C4T approach was shown to be a versatile approach for demonstrating cell-cell interactions as well as engineering cell-cell communication.

**Fig. 1.**
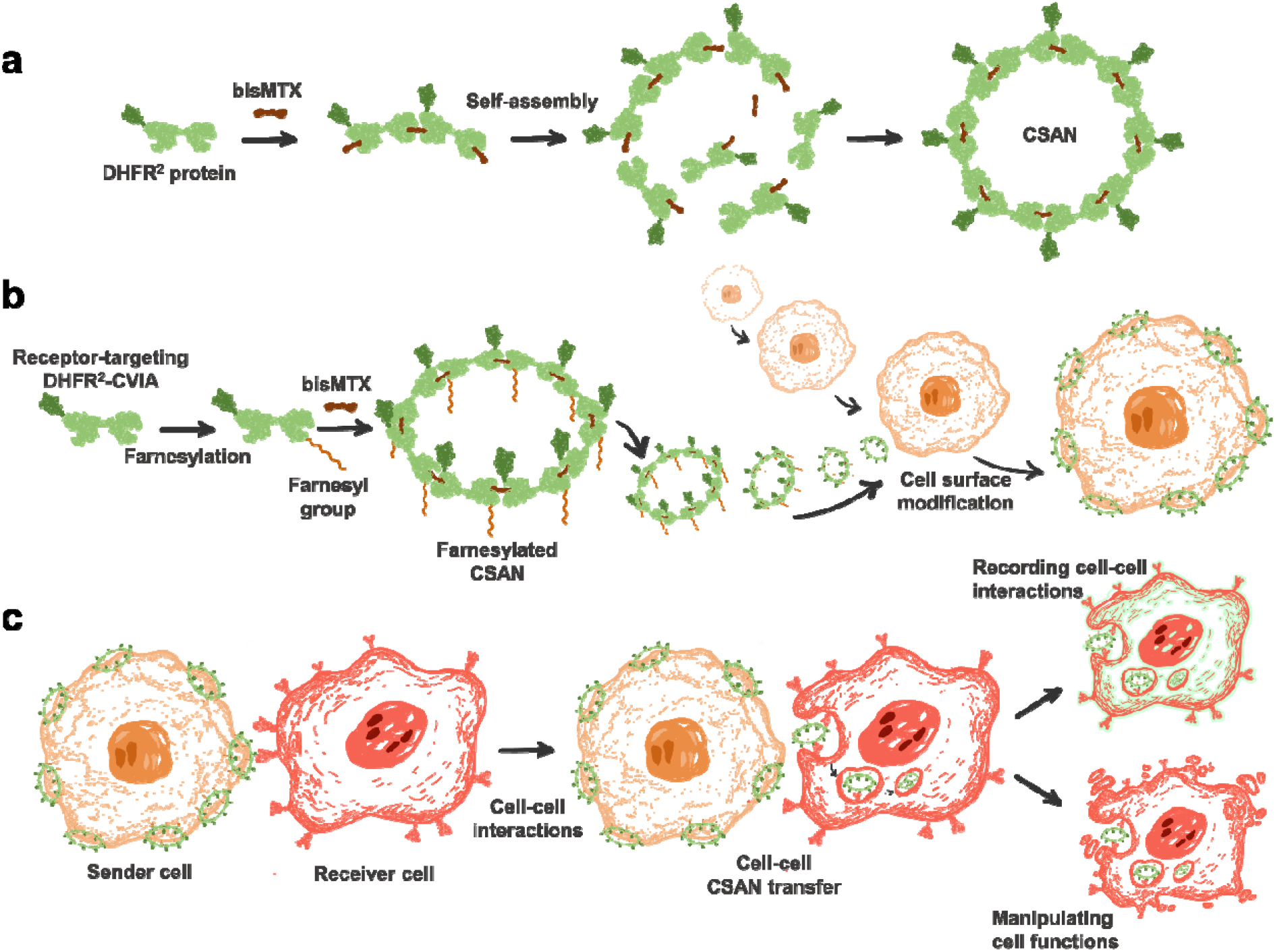
Schematic illustration of CSAN-assisted cell-cell cargo transfer to record cell-cell interactions and manipulate cell functions. **a** The CSAN was formed through the self-assembly of the DHFR^2^ fusion proteins by the chemical dimerizer, bisMTX. **b** The DHFR^2^ fusion protein that contains the targeting element and the C-terminal “CVIA” sequence can be farnesylated and oligomerized into nanorings for cell surface modification. **c** The surface-bound CSANs on the farnesyl-CSAN-modified cell (sender cell)can be transferred and internalized into the target cell (receiver cell) during cell-cell interactions. Functional molecules, such as dyes, oligonucleotides, and cancer drugs can be loaded to the farnesyl-CSANs as payloads for intercellular interaction-dependent delivery, therefore the CSAN-assisted cell-cell cargo transfer can serve as a versatile platform to record cell-cell interactions or manipulate cell functions.

## Results

### Preparation of the CSANs that target cell surface receptors

Our lab has previously developed multiple DHFR^2^ fusion protein constructs that target a variety of cancer-specific antigens, including EGFR, HER2, EpCAM, and CD133. The EGFR-targeting DHFR^2^ fusion protein (αEGFR-Fn3-DHFR^2^-CVIA) and the EpCAM-targeting protein (αEpCAM-Fn3-DHFR^2^- CVIA) contain the targeting domain that was previously generated based on the human tenth type III fibronectin (Fn3) scaffold and a C-terminal CVIA, which is a substrate for farnesyltransferase^32–35^. The HER2-targeting DHFR^2^ fusion protein (αHER2-afb-DHFR^2^-CVIA) carries an affibody-based (afb) targeting domain^36^, while the CD133-targeting DHFR^2^ fusion protein (αCD133-scFv-DHFR^2^) was prepared with the anti-CD133 scFv as the targeting element^37^. As a control and for formation of hybrid farnesyl-CSANs, a non-targeting DHFR^2^ fusion protein (DHFR^2^-CVIA) was also prepared. All the DHFR^2^ fusion proteins were expressed in *E. coli*, followed by purifications and then farnesylated by farnesyltransferase to prepare the farnesylated DHFR^2^ proteins (αEGFR-Fn3-DHFR^2^-Far, αHER2-afb-DHFR^2^-Far, αEpCAM-Fn3-DHFR^2^-Far, and DHFR^2^-Far), which were characterized by LC-MS (Fig. 2a and Supplementary Figs. 1-3). The DHFR^2^ fusion proteins were also non-specifically labeled with NHS-fluorescein as the fluorophore for the detection of CSANs on the surface of cells (Supplementary Fig. 4). The fluorescein-labeled farnesylated DHFR^2^ proteins were then self-assembled into the corresponding CSANs by bisMTX. The hydrodynamic diameters of the nanorings were characterized by dynamic light scattering (DLS) and the sizes of the fluorescein-labeled farnesylated CSANs were shown to be approximately 30 nm (Fig. 2b and Supplementary Fig. 5). Cryo-transmission electron microscopy (cryo-TEM) imaging analysis further confirmed the formation of nanoring structures and revealed that the sizes of the nanorings were consistent with the DLS analysis (Fig 2c and Supplementary Fig. 6). Moreover, the specificity of the CSANs to their target receptors was confirmed by flow cytometry demonstrating that the unfarnesylated receptor-targeting CSANs selectively bound to the receptor expressing target cells, while the non-targeting DHFR^2^-CVIA CSANs exhibited no observable binding. (Fig. 2d).

**Fig. 2.**
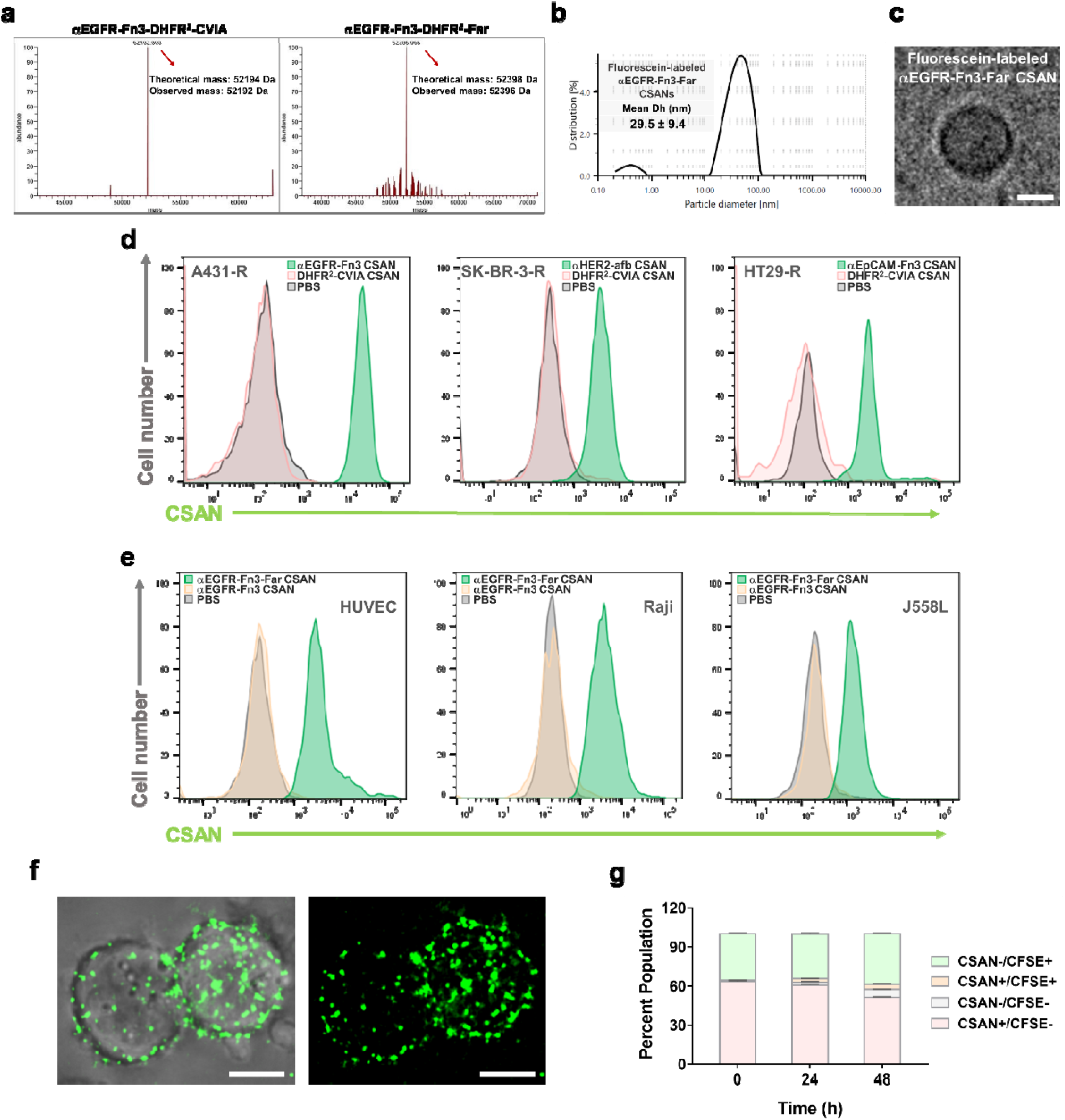
Characterizations of the CSANs and assessment of their ability for receptor binding and cell surface modification. **a** The αEGFR-Fn3-DHFR^2^-CVIA protein was efficiently farnesylated by farnesyltransferase and the proteins were characterized by LC-MS. **b** The fluorescein-labeled αEGFR-Fn3-DHFR^2^-Far protein was oligomerized into CSANs and the hydrodynamic diameter of the CSANs was measured by DLS. **c** The fluorescein-labeled αEGFR-Fn3-Far CSANs were imaged by Cryo-TEM. Scale bar, 10 nm. **d** The selectivity of the CSANs to their respective cell surface receptors was verified by flow cytometry, where the DHFR^2^ proteins were labeled with fluorescein for detection and the non-targeting DHFR^2^-CVIA CSANs served as the control. **e** Farnesyl-CSANs were shown to universally modify mammalian cell surface by flow cytometry, where the DHFR^2^ proteins were labeled with fluorescein for detection and the unfarnesylated αEGFR-Fn3 CSANs served as the control. **f** The cell surface modification of Raji cells by farnesyl-CSANs were imaged by fluorescent microscopy. Scale bar, 5 μm. **g** The stability studies of the farnesyl-CSANs on cell surface were conducted by co-culturing the CSAN-modified Raji cells and the CFSE-stained Raji cells at a 6:4 ratio in 1 mL of the medium for 48 h. The data show a negligible level of non-specific cell-cell CSAN migration over 24 h and 48 h. CSANs were labeled with DyLight-650 for detection. For **c** and **g**, data are represented as mean values ± SD (from n=3 independent experimental replicates). In some instances, small error bars are obscured by the symbols denoting the mean value. Source data are provided as a Source Data file.

### Farnesylated CSANs serve as a universal system for cell surface modifications

The farnesyl groups incorporated into the farnesylated CSANs interact with the lipid bilayer of the plasma membrane through thermodynamically favored hydrophobic insertion, thus anchoring the CSANs onto the cell surface. Previously, we have demonstrated that the prenylated CSANs are able to universally modify mammalian cell surface^19^. Farnesyl-CSANs were shown to efficiently modify primary human endothelial cells (HUVECs), human lymphoblastoid cells (Raji), and mouse myeloma cells (J558L), confirmed their ability to modify cell surfaces across species and tissue/cell types (Fig. 2e). Consistent with our previous observation that prenylated CSANs preferentially insert into lipid rafts, imaging of the modified cell surfaces by fluorescent microscopy revealed the characteristic semi-discrete localizations of the farnesyl-CSANs on the cell membranes^19^ (Fig. 2f).

Previously, due to their multivalency, farnesyl-CSANs were shown to stably modify cell surfaces (T_1/2_ > 3 days)^19^. To characterize the potential non-specific transfer of the farnesyl-CSANs to adjacent unmodified cells, the Raji cells were modified with farnesyl-CSANs that had been fluorescently labeled with DyLight-650. The CSAN-modified Raji cells were co-cultured with unmodified CFSE-labelled Raji cells, followed by incubation for 0-48 hours. Every 24 hours, aliquots of cells were analyzed by flow cytometry to measure the percentage of single-labelled and double-labelled cell populations. The amount of farnesyl-CSANs transfer was assessed by determining the percentage of CSAN^+^/CFSE^+^ proportion by flow cytometry. Over a 48-h period less than 5% of the Raji cells were shown to be CSAN^+^/CFSE^+^; thus, negligible non-specific cell-cell transfer of the farnesyl-CSANs was observed. (Fig. 2g).

### Farnesylated CSANs specifically transfer from the sender cells to the receiver cells during cell-cell interactions

To investigate if the farnesylated CSANs on the cell surface can transfer to the target cells upon cell-cell interactions, Raji cells were modified with either the fluorescein-labeled αEGFR-Fn3-Far CSANs or fluorescein-labeled non-targeting DHFR^2^-Far CSANs. The sender αEGFR-Fn3-Far-CSAN modified Raji cells were then co-cultured with the receiver EGFR^+^ A431-R cells that had been transduced to express the red fluorescent protein mKate as the marker. The co-culture conditions were carried out at 37 °C with rotation to enhance 3D interactions between the cells. After being co-cultured for 45 minutes, the cells underwent flow cytometry analysis, and the percentage of the CSAN^+^/mKate^+^ proportion was quantified to determine the extent of cell-cell transfer of the αEGFR-Fn3-Far CSANs to the A431-R cells. Within 45 minutes, 100% of the A431-R cells displayed the fluorescein signal of the αEGFR-Fn3-Far CSANs, indicating that rapid transfer of the αEGFR-Fn3-Far CSANs from modified Raji cells to the EGFR^+^ A431-R cells had occurred. No detectable farnesyl-CSAN transfer to the A431-R cells was observed with Raji cells modified with non-targeted farnesyl-CSANs (Fig. 3a,b). To verify the specificity of the transfer, an increasing amount of non-farnesylated αEGFR-Fn3 CSANs (0-5000 nM) was added to the A431-R cells and shown to block transfer of αEGFR-Fn3-Far CSANs from the CSAN-modified Raji cells. (Fig. 3f,g). Moreover, no significant transfer was observed from the αEGFR-Fn3-Far-CSAN modified Raji cells to EGFR^-^ MDA-MB-453-R (Supplementary Fig. 7). Taken together, these results are consistent with the specific transfer of αEGFR-Fn3-Far CSANs from sender cells to EGFR^+^ receiver cells.

**Fig. 3.**
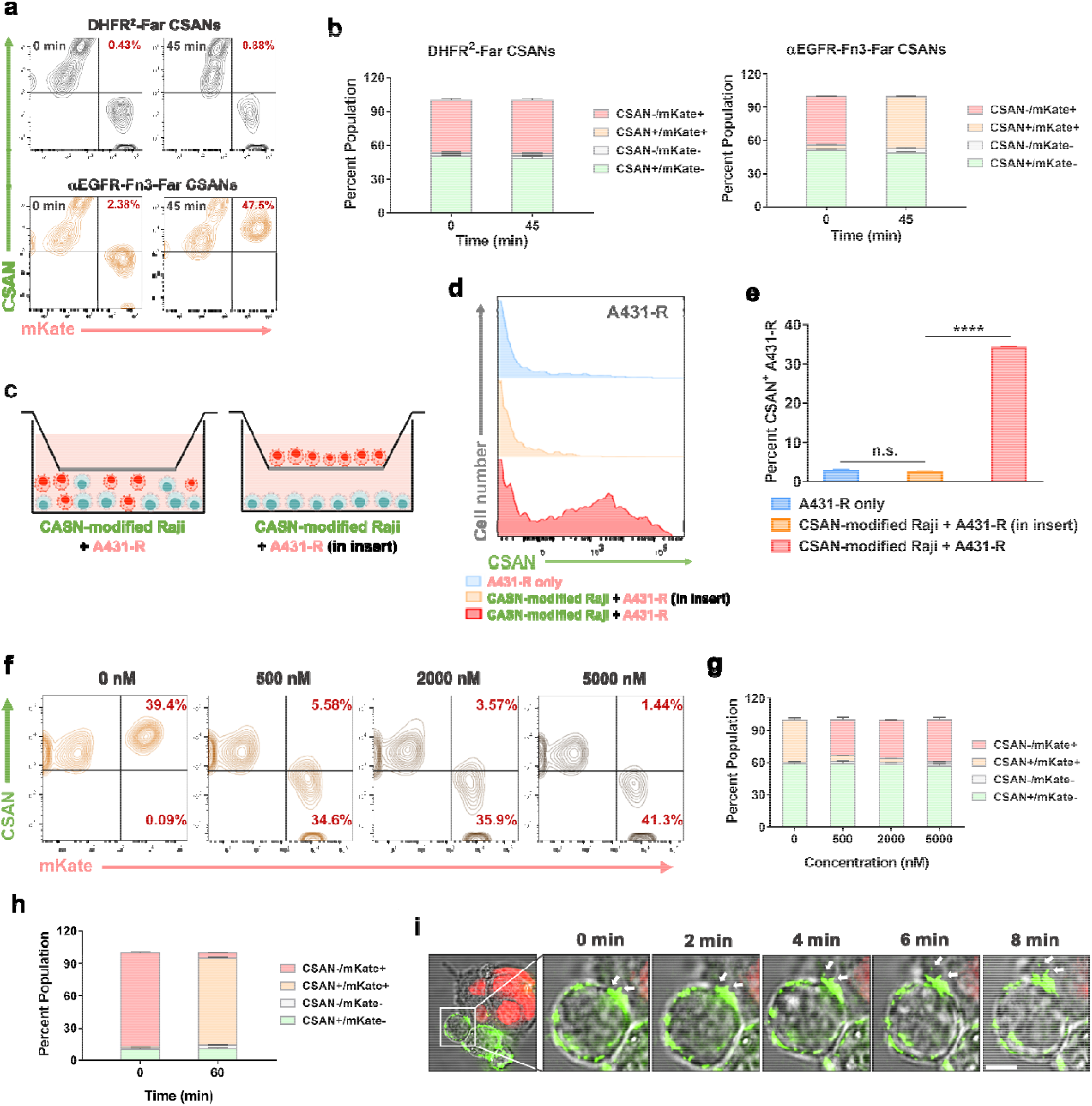
The cell-cell CSAN transfer is dependent on ligand-receptor binding and cell-cell physical contacts. **a** The representative flow cytometry plots demonstrate the proportions of cells during the co-culture of the CSAN-modified Raji cells and the A431-R cells. The percentage of CSAN^+^/mKate^+^ double-positive cell population is shown in the plot and indicates the CSAN transfer from Raji cells to A431-R cells. The quantitative data of this flow cytometry study is presented in **b** the bar graphs of percent cell populations in the cell-cell CSAN transfer study for the non-targeting DHFR^2^-Far CSANs and αEGFR-Fn3-Far CSANs. **c** Schematic illustration of the transwell assay, where the transwell inserts were used to separate the CSAN-modified Raji cells from the A431-R cells during the co-culture. **d** For the transwell assay, the A431-R cells were co-cultured with the CSAN-modified Raji cells at a 1:1 ratio with or without the transwell inserts at 37 °C for 30 min and analyzed by flow cytometry. The representative flow cytometry histograms of the transwell assay indicate the cell-cell CSAN transfer is dependent on cell-cell physical contacts. The quantitative flow cytometry data of this transwell assay is presented in **e. f** The representative flow cytometry plots of the competition assay demonstrate CSAN transfer is dependent on receptor binding. The quantitative flow cytometry data of this competition assay is presented in the **g. h** The flow cytometry study of CSAN transfer shows that the CSAN-modified Raji cells are capable of labeling multiple folds of A431-R cells through cell-cell CSAN transfer. **i** The time-lapse images show the process of the CSAN transfer from Raji cells to A431-R cells. The A431-R cells expressed red fluorescent mKate protein in the nucleus as a marker and CSANs are shown in green. The arrows highlight the punctuate spots of the CSANs. Scale bar, 5 μm. For **b, e, g, h**, data are represented as mean values ± SD (from n=3 independent experimental replicates). In some instances, small error bars are obscured by the symbols denoting the mean value. Significance in **g** was tested using a two-tailed, unpaired t-test. And is indicated as ****P < 0.0001. Source data are provided as a Source Data file.

To probe whether direct cell-cell contact is a requirement for intercellular CSAN transfer, a transwell assay with the transwell insert containing a porous membrane was carried out in which Raji cells modified with αEGFR-Fn3-Far CSANs were placed in the upper compartment, while A431-R cells were cultured in the lower compartment (Fig. 3c). As shown by flow cytometry data, for the A431-R receiver cells co-cultured together with the αEGFR-Fn3-Far-CSAN modified Raji cells, a significant increase in fluorescein intensity was observed in 30 min as expected. In contrast, when placed in the transwells, no significant fluorescein intensity was observed to have been transferred from the αEGFR-Fn3-Far-CSAN modified Raji cells to the A431-R cells (Fig. 3d,e). Rapid transfer within minutes could be observed by live-cell imaging microscopy from sender cells modified with the fluorescein labeled αEGFR-Fn3-Far CSANs to EGFR^+^ A431-R cells, followed by internalization. (Fig. 3i, Supplementary Video). Taken together, these results are consistent with the previously observed high stability of the farnesyl-CSANs on cell membranes and the need for direct physical contact between the sender cells and receiver cells for farnesyl-CSAN cargo transfer.

### Cell-cell cargo transfer and interaction are dependent on sender cell to receiver cell ratio and farnesyl-CSANs concentration

Given that, even under conditions in which transfer of the farnesyl-CSANs has resulted in 100% of the receiver cells being labelled, greater than 95% of the sender cells remain labelled with farnesyl-CSANs; thus, sender cells could in principle label multiple cells. To assess the ability of sender cells to transfer farnesyl-CSANs to multiple receiver cells, Raji cells were modified with αEGFR-Fn3-Far CSANs and co-cultured with A431-R cells in a 1:9 sender: receiver ratio, followed by flow cytometry analysis. After co-culture for one hour, CSANs were transferred from the sender cells to nearly 100% of the receiver cells without a significant decrease in the percentage of CSAN^+^ sender cells (Fig. 3h), indicating that the sender cells were able to transfer the αEGFR-Fn3-Far CSANs to multiple receiver cells. In addition, when the amount of cargo transfer over time was compared at 4 °C and 37 °C, significantly less (6-fold) αEGFR-Fn3-Far CSANs were found to have been transferred to the receiver cells indicating that the targeted farnesyl-CSANs transfer is at least partially dependent on EGFR-based endocytosis (Supplementary Fig. 8).

Previously, targeted farnesylated CSANs were shown to mediate reversible cell-cell interactions in a concentration-dependent manner^19^. To characterize the targeted farnesylated CSANs concentration dependence on the stability of induced cell-cell interactions, Raji cells were modified with variable concentrations (0-2.5 μM) of αEGFR-Fn3-Far CSANs, followed by co-culture with A431-R cells and analysis with flow cytometry. The αEGFR-Fn3-Far CSANs did not induce significant cell-cell interactions at low CSAN concentrations (< 1 μM). Nevertheless, efficient and maximal αEGFR-Fn3-Far CSANs cargo transfer was observed at similar minimal CSAN concentrations (< 1 μM) (Supplementary Fig. 9).

### The kinetic of CSAN transfer is modulated by receptor number and receptor internalization rate

Cellular receptors undergo differential rates of internalization and membrane expression. Consequently, given that αEGFR-Fn3-Far CSANs on sender cells undergoes receiver cell cargo transfer by initial binding to EGFR followed by internalization, the amount of cell-cell cargo transfer of targeted farnesyl-CSANs with time should be dependent on the rate of receptor internalization and the level of membrane expression. The cellular receptors EGFR, HER2 and EpCAM have been shown to have significantly different rates of internalization upon ligand binding (EGFR > HER2 > EpCAM)^38–40^. In addition, the expression levels of these receptors can vary drastically among cell types. Therefore, the rate of αEGFR-Fn3-Far CSANs cargo transfer and dependence of receptor cellular expression levels was compared to αHER2-afb-Far CSANs and αEpCAM-Fn3-Far CSANs. In each case we chose targeting ligands with similar affinities for their target receptors^19,34^. Both αHER2-afb-Far CSANs and αEpCAM-Fn3-Far CSANs were prepared, labelled with fluorescein and used to modify the surfaces of Raji cells. Similar numbers of Raji cells modified with similar amounts of targeted farnesyl-CSANs were co-cultured with appropriate receiver cells expressing either different levels of EGFR expression (A431-R cells, high EGFR expression; and MDA-MB-231-R cells, low EGFR expression) or similar levels of either HER2 (SK-BR-3-R) or EpCAM (HT29-R cells) (Table 1). Consistent with the 25-fold difference in EGFR expression, the time necessary for 50% of the A431-R cells receiver cells to undergo labelling with EGFR-Fn3-Far CSANs (T_50%_) from the modified Raji cells was found to be 23-fold faster than for MDA-MB-231-R cells (Fig. 4a-c,f and Table 1). Nevertheless, when normalized to EGFR expression (Table 1, T50% ^X^/T50% ^A431-R^ • NR #), no significant difference in the transferability of the αEGFR E1-Far-CSANs was observed, thus, indicating that the amount of farnesyl-CSAN cargo transfer is dependent on the level of EGFR expression and that the rate of EGFR internalization is similar for the two cell lines.

**Table 1.** Transfer kinetics of different CSANs for different receiver cell lines

|  |  |  |  |  |
| --- | --- | --- | --- | --- |
| **Receiver Cell Type** | **A431-R**<br>**(EGFR+)** | **MDA-MB-**<br>**231-R**<br>**(EGFR+)** | **SK-BR-3-R**<br>**HER2+** | **HT-29-R**<br>**EpCAM+** |

| Targeted Farnesyl CSANs | αEGFR-Fn3-Far CSAN | αEGFR-Fn3-Far CSAN | αHER2-afb-Far CSAN | αEpCAM-Fn3-Far CSAN |
| --- | --- | --- | --- | --- |
| Receptor No. | 17,212,212 ± 895,286 | 677,707 ± 15,973 | 4,328,469 ± 2,116,328 | 3,624,282 ± 16,899 |
| Normalized Receptor #<sup>a</sup> (NR#) | 1.0 | 0.0394 | 0.251 | 0.211 |
| $T_{50\%}^b$ (min) | 7 | 162 | 264 | 802 |
| $T_{50\%}^X / T_{50\%}^{A431-R} \bullet NR \#$ | 1.0 | 0.912 | 9.47 | 24.2 |

<sup>a</sup> Normalized Receptor # (NR#) = Cell Receptor #/A431-R EGFR #
<sup>b</sup> $T_{50\%}$ , the time required for 50% of the receiver cells to undergo labelling with targeted farnesyl-CSANs.

**Fig. 4.**
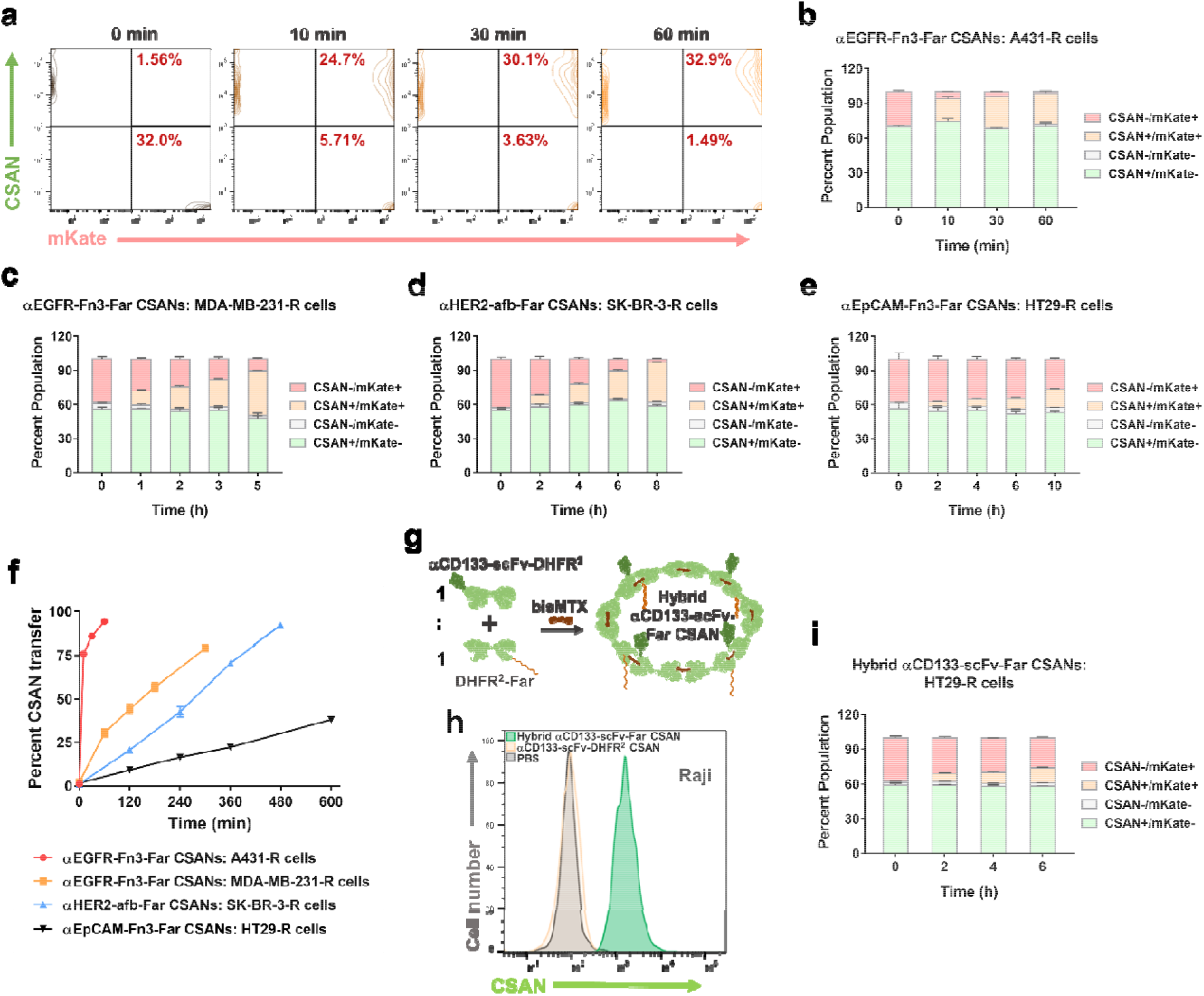
Kinetics study of the cell-cell CSAN transfer for different cell lines and receptors. **a** The representative flow cytometry plots of the kinetics study for the αEGFR-Fn3-Far CSANs transferring from Raji cells to A431-R cells. The quantitative flow cytometry data of this kinetics study was presented in **b. c** The kinetics study for the αEGFR-Fn3-Far CSANs transferring from Raji cells to MDA-MB-231-R cells. **d** The kinetics study for the αHER2-afb-Far CSANs transferring from Raji cells to SK-BR-3-R cells. **e** The kinetics study for the αEpCAM-Fn3-Far CSANs transferring from Raji cells to HT29-R cells. **f** The plot of transfer kinetics of the CSANs for different target cells. The percent CSAN transfer is determined by the percentage of the CSAN^+/^mKate^+^ cell proportion out of the mKate^+^ receiver cell proportion. **g** Schematic illustration of the formation of the hybrid αCD133-scFv-far CSANs. **h** The hybrid αCD133-scFv-far CSANs were shown to adequately modify Raji cell surface. **i** The kinetics study for the hybrid αCD133-scFv-far CSANs transferring from Raji cells to HT29-R cells. For **b-f** and **i**, data are represented as mean values ± SD (from n=3 independent experimental replicates). In some instances, small error bars are obscured by the symbols denoting the mean value. Source data are provided as a Source Data file.

When cargo transfer of αEGFR-Fn3-Far CSANs was compared to αHER2-afb-Far CSANs, αEGFR-Fn3-Far CSANs labelling (T_50%_) of A431-R (EGFR^+^) cells was found to be 38-fold greater than αHER2-afb-Far CSANs cargo transfer to SK-BR-3-R (HER2^+^) cells, despite only a 4-fold lower expression of HER2 than EGFR. Consistent with the approximately 8-fold difference in the rate of HER2 internalization relative to the rate of EGFR internalization^39,40^, the T_50%_^X^/T_50%_^A431-R^ • NR # value for αHER2-afb-Far CSANs and SK-BR-3-R (HER2^+^) cells was found to be 9.47-fold greater than observed for αEGFR-Fn3-Far CSANs transferred to A431-R (EGFR^+^) cells. Similarly, the T_50%_^X^/T_50%_^A431-R^ • NR # value for αEpCAM-Fn3-Far CSANs was found to be approximately 24-fold and 2.5-fold greater than αEGFR-Fn3-Far CSANs labelling (T_50%_) of A431-R (EGFR^+^) cells and αHER2-afb-Far CSANs for SK-BR-3-R (HER2^+^) cells, respectively. Thus, when normalized to the level of receptor expression, despite the difference in cell type, the cargo transferability of the targeted farnesyl-CSANs correlates with the rate of receptor internalization.

### Modular design of CSANs facilitates use of single chain antibody targeting ligands

In general, the use of the C-terminal CVIA bio-conjugation tag requires that a targeting ligand fused to the N-terminus, lack disulfides. In particular, the incorporation of the widely used single chain antibody frame work in the presence of additional cysteines in the fusion protein can lead to difficulties to manipulate aggregates, even after undergoing careful refolding or expression in disulfide isomerase expressing bacterial strains^41,42^. Previously, we demonstrated that farnesyl-and geranylgeranyl-CSANs displaying variable amounts of prenylation could be prepared by the self-assembly of prenylated monomers with unprenylated targeting monomers^19^. Indeed, the stability of farnesyl-CSANs composed of an average of four farnesylated DHFR^2^ monomers was sufficient for maximum stable binding to cell surfaces^19^.

CD133 is a transmembrane protein that has been found to be associated with neural and haematopoietic stem cells, as well as cancer stem-like cells (CSC)^43,44^. Although, stem cells and CSCs have been shown to be resistant to chemotherapeutics, potent immunotoxins targeting CD133 are able to be internalized and kill tumor cells expressing CD133^45–47^. Nevertheless, the potential for sender cells to target CD133 expressing receiver cells for cargo transfer has, to our knowledge, not been reported. Consequently, to investigate the potential for utilizing farnesyl-CSANs incorporating disulfides-containing targeting ligands for cell-cell cargo transfer, the hybrid αCD133-scFv-far CSANs were self-assembled by mixing a 1:1 ratio of αCD133 scFv-DHFR^2^ and DHFR^2^-Far monomers in the presence of bisMTX (Fig. 4g and Supplementary Fig. 10). Raji cells were modified with the hybrid αCD133-scFv-far CSANs (Fig. 4h), incubated with CD133^+^ HT29-R cells and the transfer of the fluorescein-labelled CSANs monitored over time by flow cytometry (Fig. 4 f,i and Supplementary Fig. 11). The T_50%_ values for aEGFR-Fn3-Far CSANs labelling of A431-R (EGFR^+^) cells was found to be 78-fold greater than hybrid aCD133-scFv-Far CSANs cargo transfer to HT29-R (CD133^+^) cells. Unfortunately, an aCD133 monoclonal antibody that binds to both glycosylated and non-glycosylated CD133 is not available, precluding our ability to accurately measure the amount of CD133 on HT-29-R cells and thus directly compare the internalization behavior of CD133 with other receptors. Nevertheless, the C4T results with hybrid aCD133-scFv-Far CSANs are consistent with prior studies demonstrating that the aCD133-scFv could be employed for drug delivery^45–48^.Thus, hybrid targeted farnesyl-CSANs and scFvs can be used to carryout C4T.

### Dependence of cell-cell cargo transfer on sender and receiver cell type

Having investigated the ability of Raji cells to serve as sender cells, we chose to investigate the ability of different sender cells and receiver cells to carry out cargo transfer. Cytotoxic T-lymphocytes (CD8^+^ T cells) and Natural Killer (NK) cells were modified with fluorescein-labeled αEGFR-Fn3-far CSANs or control fluorescein-labeled DHFR^2^-Far CSANs and their ability to facilitate cargo transfer to A431-R (EGFR^+^) cells over 30 min was determined by flow cytometry and fluorescent microscope (Fig. 5a-f). While no significant transfer of non-targeted farnesyl-CSANs to A431-R was observed for either CD8^+^ T cells or NK cells, significant cargo transfer was observed for αEGFR-Fn3-far CSANs to the target cells. Interestingly, although the sizes of CD8^+^ T cells or NK cells are very similar, over twice as much αEGFR-Fn3-far CSANs was found to be transferred by NK cells relative to CD8^+^ T cells. Consistent with these findings, internalized fluorescently labelled punctate spots were observed in the receiver A431-R cells by fluorescent microscopy. The punctate spots, and thus labelled αEGFR-Fn3-far CSANs, could be observed throughout the cell and even near or associated within the nucleus.

To examine the feasibility of farnesyl-CSAN-based cargo transfer with non-lymphocytic cells, human vascular endothelial cells (HUVEC) were modified with fluoresceine labelled αHER2-afb-far CSANs or non-binding control farnesyl-CSANs and the amount of cargo transfer to SK-BR-3-R (HER2^+^) cells determined by flow cytometry and fluorescence microscopy. HUVEC cells were indeed shown to serve as sender cells for αHER2-afb-far CSANs transferring to SK-BR-3-R cells, with multiple fluorescein-labelled punctates observable through the cell. (Figure 5 j-1) The potential for phagocytotic cells, such as macrophages, to serve as receiver cells was also examined. Mouse myeloma cell line, J558L, were modified with fluorescein-labelled αEGFR-Fn3-far CSANs and incubated with mouse macrophage cells, RAW 264.7 (EGFR^+^) for 30 min. Macrophages have been shown to express EGFR, which has been shown to play a critical role in their response to pathogens^49^. Significant amounts of cargo transfer of the labelled farnesyl-CSANs were observed by flow cytometry with green fluorescent punctates observable by fluorescent microscopy throughout the RAW 264.7 receiver cells including the nucleus. (Figure 5 g-i) Previously, EGFR-targeted CSANs were shown to be endocytosed by macrophage RAW 264.7 cells^50^, in addition, non-EGFR targeted CSANs were shown to undergo uptake by the common scavenger receptor-1 (SR-1)^51^. Interestingly, non-specific cargo-transfer was not observed by the non-targeted Farnesyl-CSANS to RAW 264.7. Taken together, these examples, indicate that farnesyl-CSANs can be used to label a variety of sender cell types in which a variety of receiver cells expressing the target receptors, in this case EGFR and HER2, can carry out cargo transfer by endocytosis.

**Fig. 5.**
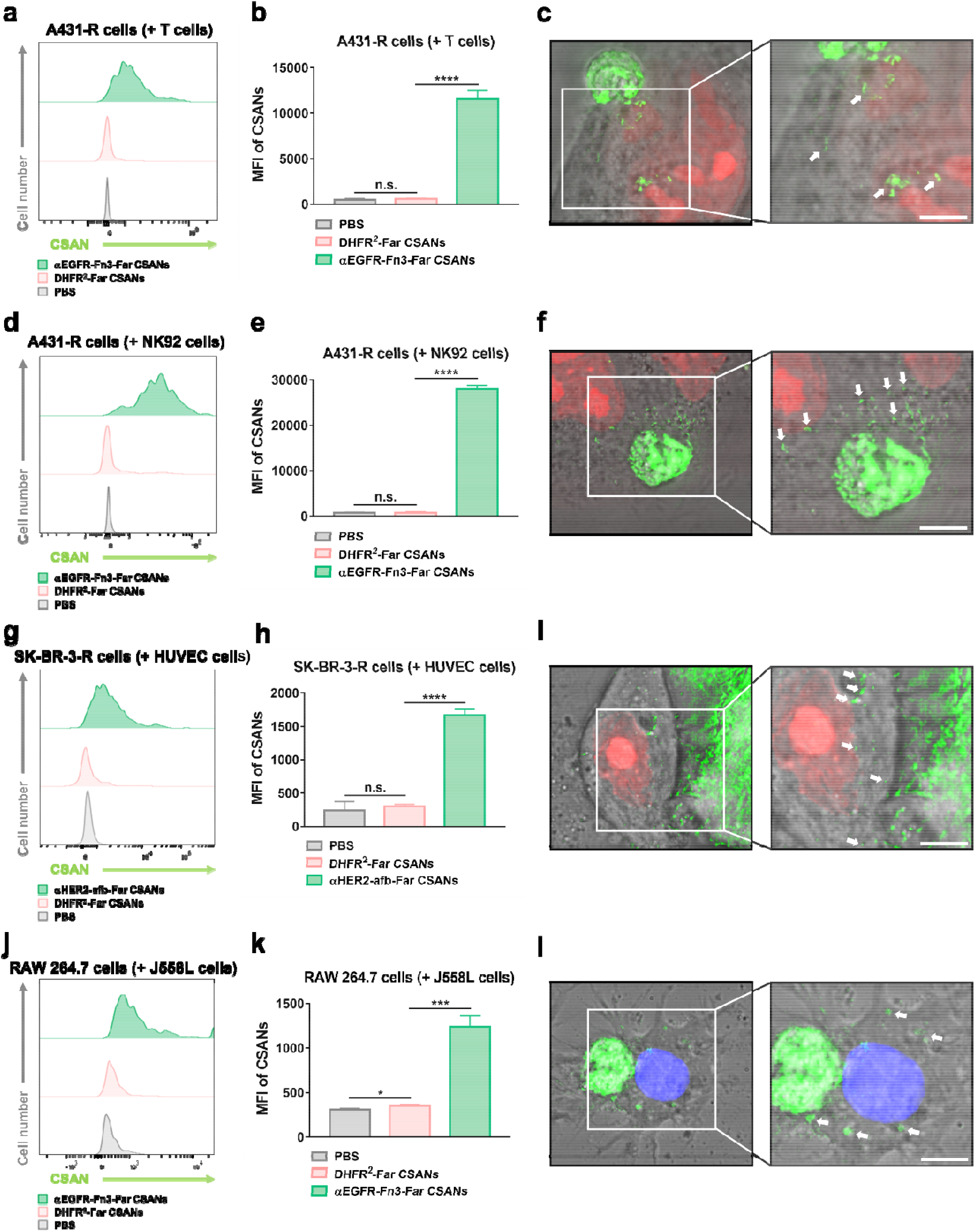
Recording cell-cell interactions by CSAN-assisted cell-cell cargo transfer. **a** The representative flow cytometry histograms show the CSANs on the A431-R receiver cells after the co-culture with CSAN-modified T cells. The quantitative flow cytometry data are presented in **b. c** The T-cancer cell-cell interactions were recorded by fluorescence microscopy with the C4T approach. **d** The representative flow cytometry histograms show the CSANs on the A431-R receiver cells after the co-culture with CSAN-modified NK92 cells. The quantitative flow cytometry data are presented in **e. f** The NK-cancer cell-cell interactions were recorded by fluorescence microscopy with the C4T approach. **g** The representative flow cytometry histograms show the CSANs on the RAW 264.7 receiver cells after the co-culture with CSAN-modified J558L cells. The quantitative flow cytometry data are presented in **h. i** The myeloma-macrophage cell-cell interactions were recorded by fluorescence microscopy with the C4T approach. **j** The representative flow cytometry histograms show the CSANs on the SK-BR-3-R receiver cells after the co-culture with CSAN-modified HUVEC cells. The quantitative flow cytometry data are presented in **k. l** The endothelial-cancer cell-cell interactions were recorded by fluorescence microscopy with the C4T approach. In the flow cytometry study and the imaging study, A431-R cells and SK-BR-3-R cells expressed red fluorescent mKate protein in the nucleus as the marker, and RAW 264.7 cells were stained with the Hoechst dye and shown in blue. All CSANs are shown in green. For **c, f, i, l**, the arrows highlight the punctuates of the CSANs. Scale bars, 5 μm. For **b, e, h, k**, data are represented as mean values ± SD (from n=3 independent experimental replicates). In some instances, small error bars are obscured by the symbols denoting the mean value. Significance in these plots was tested using a two-tailed, unpaired t-test and is indicated as *P < 0.05, ***P < 0.001 and ****P < 0.0001. Source data are provided as a Source Data file.

### Interaction-dependent delivery of an anti-cancer drug

Having demonstrated that targeted farnesyl-CSANs are able to carry out cargo transfer, their potential to selectively deliver, not only dyes, but biologically active agents, such as drugs was investigated. Since farnesyltransferase is a promiscuous enzyme that can employ isoprenoids other than native farnesyl diphosphate as substrates^52–54^. The αEGFR-Fn3-DHFR^2^-CVIA monomer was farnesylated with farnesyl-azide diphosphate (C10-N_3_-OPP) to prepare αEGFR-Fn3-DHFR^2^-N_3_, followed by click conjugation to dibenzocyclooctyne covalently attached to valine-citrulline-p-aminobenzoyloxycarbonyl-monomethyl auristatin E conjugated through a PEG linker (DBCO-PEG_4_-VC-PAB-MMAE) (Supplementary Fig. 12). VC-PAB-MMAE incorporates a cathepsin sensitive valine-citrulline-PAB self-immolative linker and the potent anti-mitotic agent, MMAE. The drug linker combination has been extensively employed for the development of antibody drug conjugates, including several FDA-approved ADCs^55^. The resultant protein-drug conjugate, αEGFR-Fn3-DHFR^2^-MMAE, was self-assembled in the presence of αEGFR-Fn3-DHFR^2^-Far at a 1:3 ratio, respectively, into hybrid αEGFR-Fn3-Far-MMAE CSANs (Fig. 6a and Supplementary Fig. 13). Raji cells were modified with hybrid αEGFR-Fn3-Far-MMAE CSANs and found to have no significant effect on cell viability, thus they do not undergo internalization by the sender cells. Sender Raji cells were modified with fluorescently labelled αEGFR-Fn3-Far-MMAE CSANs, followed by the co-culture with A431-R cells. Within 30 mins, cargo-transfer of the hybrid αEGFR-Fn3-Far-MMAE CSANs could be observed by fluorescent microscopy with fluorescent punctates observable throughout the cell, including the nucleus (Fig. 6b,c and Supplementary Fig. 14).

**Fig. 6.**
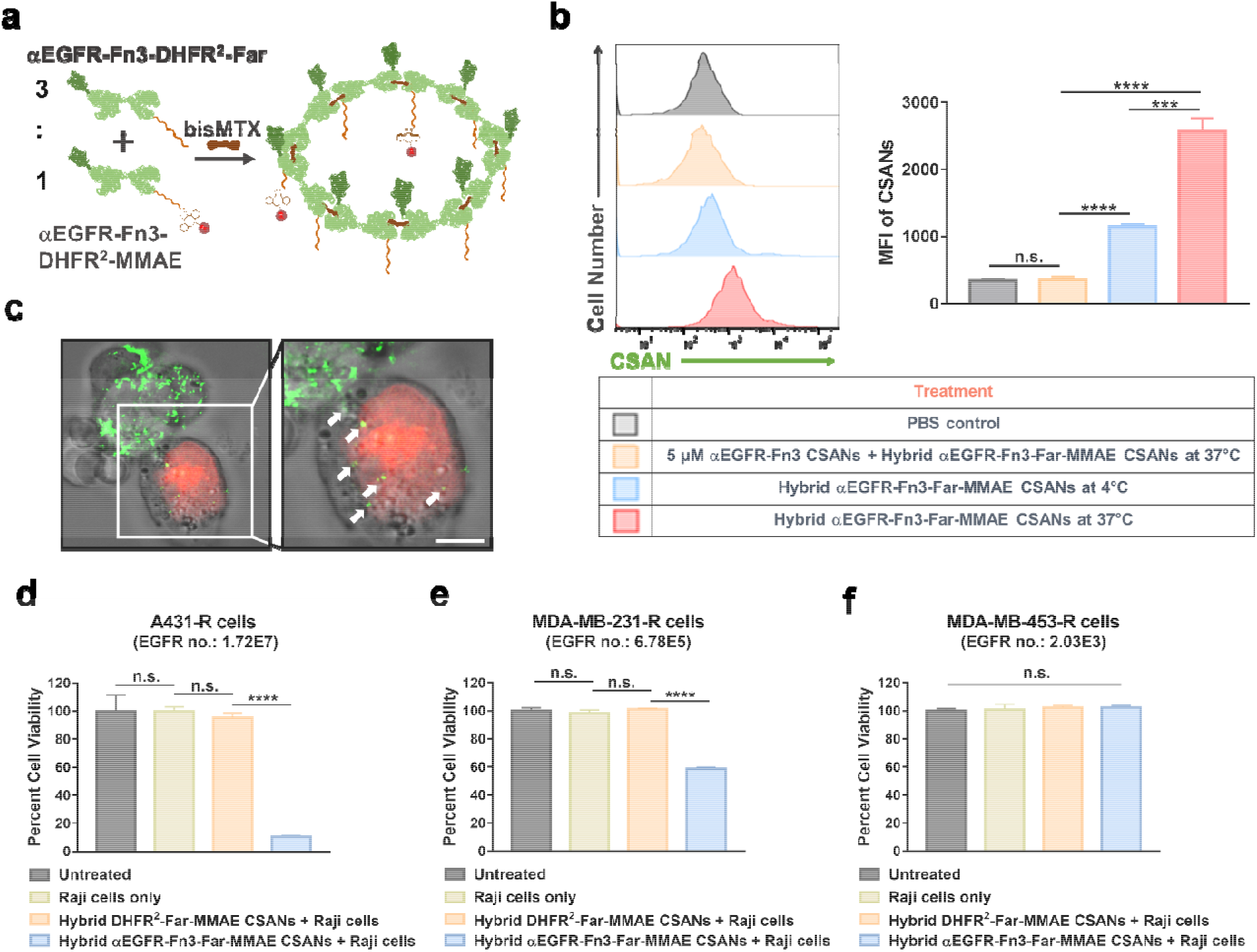
Study of the cell-cell interaction-dependent delivery of the anti-cancer drug by the C4T approach. **a** Schematic illustration of the formation of the hybrid αEGFR-Fn3-Far-MMAE CSANs. **b** The flow cytometry study of the cell-cell interaction-dependent delivery of MMAE by the C4T approach. The αEGFR-Fn3-DHFR^2^-MMAE protein conjugate was labeled with fluorescein for detection. **c** The fluorescent images indicate the transfer of the hybrid αEGFR-Fn3-Far-MMAE CSANs from Raji cells to A431-R cells. The A431-R cells expressed red fluorescent mKate protein in the nucleus as a marker and CSANs are shown in green. Scale bar, 5 μm. The cytotoxicity of the cell-cell interaction-dependent delivery of MMAE by the C4T approach was studied using IncuCyte for **d** A431-R cells, **e** MDA-MB-231-R cells, and **f** MDA-MB-453-R cells, where the CSAN-modified Raji cells were co-cultured the target cancer cells at a 3:1 ratio in the plate at 37 °C for 2 hours, followed by medium exchange to remove the Raji cells. The cancer cells were returned to the IncuCyte and cultured for 4 days to quantify cell viability. For **b** and **d-f**, data are represented as mean values ± SD (from n=3 independent experimental replicates). In some instances, small error bars are obscured by the symbols denoting the mean value. Significance in **b** and **d-f** was tested using a two-tailed, unpaired t-test. And is indicated as ****P < 0.0001 and ***P < 0.001. Source data are provided as a Source Data file.

To assess the potential for the hybrid αEGFR-Fn3-Far-MMAE CSANs to act as drug delivery vehicles, sender Raji cells were modified with hybrid αEGFR-Fn3-Far-MMAE CSANs, followed by co-culturing with A431-R (17×10^6^ EGFR per cell) cells for 2 h. The CSAN-modified Raji cells were then removed from the co-culture and the A431-R cells cultured for 96 h, followed by the cell viability quantification. Co-culturing with Raji cells modified with the αEGFR-Fn3-Far-MMAE CSANs resulted in potent cytotoxicity against A431-R cells, while the unmodified Raji cells or Raji cells modified with non-targeting hybrid DHFR^2^-Far-MMAE CSANs failed to exert any significant cytotoxicity toward the receiver cells (Fig. 6d). To assess the role of EGFR expression on the cargo transfer induced cytotoxicity by the hybrid αEGFR-Fn3-Far-MMAE CSANs, a similar cytotoxicity study was carried out with MDA-MB-231-R cells and MDA-MB-435-R, which express 25-fold and 8500-fold less EGFR than A431-R cells, respectively. Clearly, the expression levels of EGFR on the receiver cells impacted the ability of the sender Raji cells modified with hybrid αEGFR-Fn3-Far-MMAE CSANs to carry out cargo transfer induced cytotoxicity, since a 6-fold reduction in cytotoxicity was observed for MDA-MB-231-R receiver cells, while no significant cytotoxicity was observed for the MDA-MB-435-R cells (Fig. 6e,f).

### Interaction-dependent delivery of functional oligonucleotides

Given that targeted Farnesyl-CSANs are able to facilitate cargo transfer of small molecules, we investigated their ability to carry out macromolecular delivery by assessing oligonucleotide transfer. αEGFR-Fn3-DHFR^2^-N3 was treated with a DNA oligonucleotides modified with a DBCO group at the 5’-terminus and AlexaFluor-488 fluorophore at the 3’-terminus (Supplementary Fig. 15). The resultant protein-oligonucleotides conjugate, αEGFR-Fn3-DHFR^2^-ssDNA-AF488, was self-assembled in the presence of αEGFR-Fn3-DHFR^2^-Far at a 1:3 ratio, respectively, into hybrid αEGFR-Fn3-Far-ssDNA-AF488 CSANs (Fig. 7a and Supplementary Fig. 16). The plasma membranes of the sender Raji cells were modified with the hybrid CSANs, and 5-fold more of the fluorescently labeled hybrid αEGFR-Fn3-Far-ssDNA-AF488 CSANs was observed to have been transferred to the receiver A431-R cells at 37 °C than 4 °C (Fig. 7b).

**Fig. 7.**
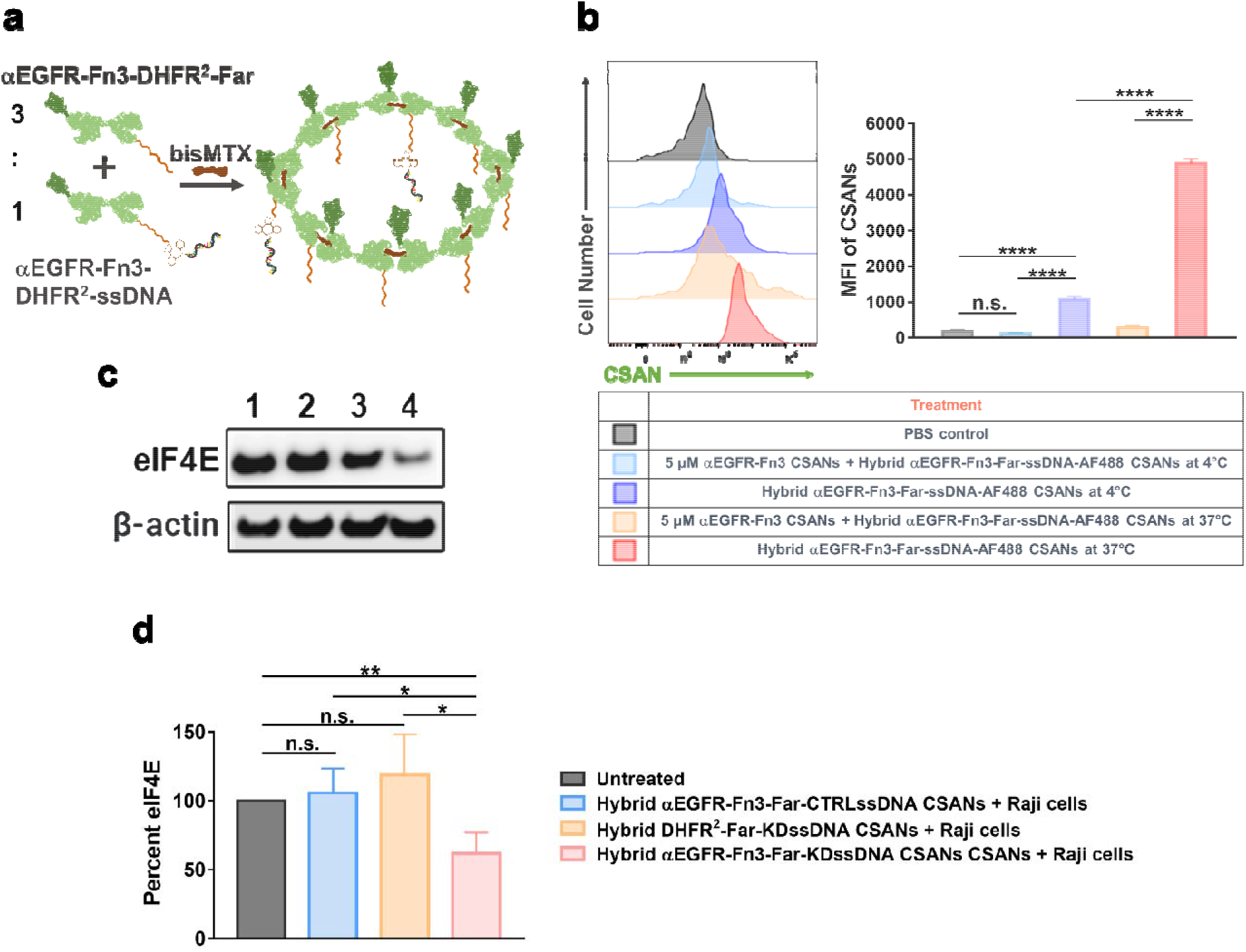
Study of the cell-cell interaction-dependent delivery of oligonucleotides by the C4T approach. **a** Schematic illustration of the formation of the hybrid αEGFR-Fn3-Far-ssDNA CSANs. **b** The flow cytometry study of the cell-cell interaction-dependent delivery of ssDNA by the C4T approach. The αEGFR-Fn3-Far-ssDNA-AF488 protein conjugate has an AlexaFluor-488 dye for detection. **c** The representative western blot image shows the specific knockdown of eIF4E in MDA-MB-231 cells by the interaction-dependent delivery of the anti-eIF4E antisense ssDNA using the C4T approach. Lane 1: Raji cells modified with the hybrid DHFR^2^-Far-KDssDNA CSANs; lane 2: Raji cells modified with the hybrid αEGFR-Fn3-Far-CTRLssDNA CSANs; lane 3: untreated control; lane 4: Raji cells modified with the hybrid αEGFR-Fn3-Far-KDssDNA CSANs. **d** The quantitative western bolt data are summarized in the bar graph and are represented as mean values ± SD (from n=3 independent biological replicates). In some instances, small error bars are obscured by the symbols denoting the mean value. Significance in **b** and **d** was tested using a two-tailed, unpaired t-test. And is indicated as *P < 0.05, **P < 0.01, and ****P < 0.0001. Source data are provided as a Source Data file.

Previously, both bivalent and octavalent anti-α_v_β_3_ CSANs composed of DHFR^2^-cyclic-RGD monomers and bisMTX conjugated to the anti-eIF4E antisense oligonucleotides, KDssDNA, was shown to knock-down eIF4E translation in MDA-MB-231 cells by approximately 50%, when compared to anti-α_v_β_3_ CSANs composed of a scrabbled DNA oligo^56^. Consequently, given that EGFR targeting has been used for nanoparticle based delivery of a variety of nucleic acids^57^, the hybrid αEGFR-Fn3-Far-KDssDNA CSANs were prepared by oligomerizing αEGFR-Fn3-DHFR^2^-KDssDNA and αEGFR-Fn3-DHFR^2^-Far at a 1:3 ratio. Raji sender cells were modified with either the hybrid aEGFR-Fn3-Far-KDssDNA CSANs or the hybrid aEGFR-Fn3-Far-CTRLssDNA CSANs, which contained a scrambled control ssDNA (CTRLssDNA). After co-culturing of the modified sender Raji cells with MDA-MB-231 cells for 48 hours, the amount of intracellular eIF4E was quantified by western blot analysis. Compared to Raji cell modified with αEGFR-Fn3-Far-CTRLssDNA CSANs, a greater than 40% reduction in the amount of eIF4E was observed for MDA-MB-231 receiver cells co-cultured with Raji sender cells modified with the hybrid αEGFR-Fn3-Far-KDssDNA CSANs. (Fig. 7c,d). Although not optimized for farnesyl-CSAN concentration, sender cell-receiver cell ratio, conjugation linker length, and intracellular ASO release, the ability to observe significant activity for the anti-eIF4E ASO suggests that cell-cell cargo transfer may be used for functional cell based delivery of not just small molecules, but also nucleic acids.

## Discussion

Cell-cell communications is carried out by either a soluble hormone with a cell membrane receptor or the direct engagement of a membrane bound ligand with a membrane bound receptor. Of these two modes of contact, the direct contact between two different cells can result not just in receptor activation and intracellular signal transduction, but in many cases the direct transfer and internalization of the ligand from one cell membrane to the other cell^58–63^. The amount of transfer from sender cells can span from a few ligands to significant amounts of the sender cell membrane, referred to as trogocytosis. Trogocytosis has been observed between immune cells and antigen presenting cells, neurons and microglia, parasites, and endothelia cells, to name a few^58,61,62^ The unexpected induction of trogocytosis has even been observed between anti-CD19 CAR-T-cells and B-cell leukemia^64^. Recently, Tang and co-workers have pioneered the development of a cargo transfer based approach for monitoring cell-cell interactions^16^. Sender cells are genetically engineered to express GFP or a GFP analog fused to a membrane displaying domain. Cargo transfer is then facilitated by receiver cells expressing an anti-GFP nanobody fused to an internalizing membrane domain, referred to as GFP-based Touching Nexus, or G-baToN. Interestingly, the G-baToN approach has been shown to enable cell based delivery of fluorophores, proteins and nucleic acids from sender to receive cells as possible tools for monitoring cell-cell interactions^16^. Inspired by Tang et al, we investigated the potential of developing a non-genetic biomimetic trogocytosis approach for engineering cell-cell cargo transfer that could be employed with a variety of cell types, irrespective of their ability to be stably transfected, and potentially useful for targeted cell-based drug delivery, as well as cell-cell interaction monitoring.

Our laboratory has demonstrated that targeted chemically self-assembled nanorings (CSANs) can be prepared by incubation of DHFR-DHFR (DHFR^2^) fusion proteins recombinantly coupled to a targeting single chain antibody (scFv), fibronectin (Fn3), affibody (afb) or peptide^22,34,50,56^. CSANs that can covalently or non-covalently couple modified non-natural phospholipids have been shown to modify cell membranes^19,33,65^. Recently, we have demonstrated that DHFR^2^ monomers fused with a C-terminal CVIA sequence can be farnesylated or geranylgeranylated by farnesyltransferase or geranylgeranyltransferase respectively. Self-assembly of the prenylated monomers targeted to specific cell surface antigens, resulted in prenylated CSANs capable of inducing specific cell-cell interactions, such as T-cell killing of the target tumor cells^19^. Nevertheless, although the prenylated CSANs were found to stably bind to cell membranes for days, due to their multivalency, we reasoned that receptor targeted farnesyl CSANs on sender cells could undergo cargo transfer through energy dependent internalization of the targeted plasma membrane receptor.

Analysis of the temperature dependence of targeted farnesyl-CSANs cargo transfer demonstrated that cargo transfer from sender cells to receiver cells was carried out by energy dependent receptor internalization (Supplementary Fig. 8). In addition, when compared across receptors and considering that the binding affinities of the monomeric binding ligands are similar (<10-fold), the extent of cargo transfer was found to be dependent on the interplay between receptor internalization rate and level of receptor expression. For example, when comparing EGFR based cargo transfer from cell lines (A431-R vs MDA-MB-231-R) differing in EGFR expression by 25-fold, the amount of cargo transfer was found to be similarly decreased by 23-fold for the lower expressing cell line (MDA-MB-231) (**Table 1**). Nevertheless, as expected, the relative efficiency of internalization (REI) for the targeted receptor was found to be similar, indicating that the rate of EGFR internalization is independent on the cell type (**Table 1**). Similarly, when the amount of cargo transfer between two cell lines expressing similar levels of receptors, but with different internalization rates, was compared, cells expressing the faster internalizing receptor (HER2, SK-BR-3-R) accumulated a greater amount of farnesyl-CSANs than cells expressing the slower internalizing receptor (EpCAM, HT-29-R). Using EGFR expression by A431-R cells as a benchmark, the relative internalization efficiency (REI) could be determined for EGFR, HER2 and EpCAM, with EGFR being the most efficient and EpCAM the least efficient. (**Table 1**) Consistent with these findings, the rate of EGFR internalization has been found to be 8-to 10-fold faster than HER2 and > 10-fold faster than EpCAM, depending on the cellular differentiation conditions^38–40^. (**Table 1**) Consequently, targeted farnesyl CSANs cargo transfer can be used to assess the relative internalization efficiencies of cell surface receptors for targeting ligands, if the number of receptors is known and the time necessary to transfer the targeted farnesyl-CSANs to 50% of the cells is known (**Table 1**).

Given the modularity of CSANs, the ability to prepare bispecific CSANs and the observation that CSANs composed of a 4:4 ratio of DHFR^2^-Far and a targeted-DHFR^2^ stably bound cell membranes as those composed of 8 farnesyl-DHFR^2^, hybrid αCD133-scFv-Far CSANs were prepared from DHFR^2^-Far and αCD133-scFv-DHFR^2^. The hybrid αCD133-scFv-Far CSANs were shown to bind stably to the sender cell membranes and to carry out cargo transfer to CD133^+^ HT-29 R cells. Interestingly, αEpCAM-Fn3-Far CSANs and the hybrid αCD133-scFv-Far CSANs were shown to carry out similar levels of cargo transfer over a similar time frame. Thus, both farnesyl-CSANs could be used to carry out cargo transfer to the same cells over the same time frame opening the door to future studies of the effect of multi-specific targeting ligands on cargo transfer due to multispecific targeting. Accessing multi-receptor internalization, could allow for more precise cargo transfer to specific cells based on their receptor expression level and heterogeneity.

The modular nature of CSANs self-assembly suggested that delivery of conjugated payloads might also be assessable. Hybrid αEGFR-Fn3-Far-MMAE CSANs were prepared by bisMTX facilitated self-assembly of αEGFR-Fn3-DHFR^2^-MMAE and αEGFR-Fn3-DHFR^2^-Far. Sender cells loaded with the hybrid αEGFR-Fn3-Far-MMAE CSANs delivered the drug-loaded farnesyl-CSANs followed by the induction of cell death. Consistent with the finding that the efficiency of cargo transfer is dependent on receptor expression, the amount of observed cytotoxicity was found to be dependent on EGFR expression levels. Similarly, hybrid αEGFR-Fn3-Far-ssDNA CSANs were prepared from αEGFR-Fn3-DHFR^2^-ssDNA and αEGFR-Fn3-DHFR^2^-Far and found to efficiently transfer oligonucleotides from sender cells to receiver cells. In addition, if the oligonucleotide was an anti-sense oligonucleotide targeting the translation initiation factor eIF4E, significant knockdown of the amount of eIF4E was observed in the receiver cells. Importantly, the sender cells were not affected by the targeted farnesyl-CSANs cargos bound to their membranes. Thus, targeted farnesyl-CSANs could potentially be used for drug delivery to tissues that the sender cells have an affinity too. For example, the natural affinity of mesenchymal stem cells for tumor sites has been shown to enhance tumor targeting and penetration without adverse off target effects and tumor penetration challenges typically observed for nano and micro-based drug delivery carriers^66^. In addition, in principle multiple drugs or oligonucleotides could be loaded onto sender cells by simply incubating mixtures of targeted farnesyl-CSANs conjugated to specific payloads with sender cells. Importantly, unlike other nanoparticle delivery approaches, the tissue localization and biodistribution of the CSANs will likely be dependent on the tissue penetration and biodistribution of the cells and not the nanorings. In essence, regardless of what is being delivered, targeted farnesyl-CSANs based cell-cell cargo transfer can be used to non-genetically engineer synthetic cell-cell interaction tracking and communications in which chemical messages are delivered by sender cells to specific receiver cells in order to mark them or induce a desired response, be it apoptosis or alternations in signal transduction pathways. With the inherent modular nature and universal membrane binding ability of the CSANs based cell-cell cargo transfer approach, multi-targeted cargo transfer can be explored from any sender cell of choice with potentially greater cargo transfer specificity obtained to the receiver cell of choice. Studies exploring the efficacy of targeted farnesyl CSANs cargo transfer *in vivo* are underway; the results of which will be reported in due course.

## Methods

### Cell lines and cell culture

The A431, MDA-MB-231, MDA-MB-453, SK-BR-3, HT29, RAW 264.7, HUVEC, Raji, and NK-92 cells were previously purchased from the American Type Culture Collection (ATCC). The J558L cells were obtained from Dr. Bruce Walxheck. The A431, MDA-MB-231, MDA-MB-453, SK-BR-3, and HT29 cells were transfected to express the nuclear-restricted mKate2 red fluorescent protein using the IncuCyte® NucLight Lentivirus Reagents (Sartorius, Cat: 4476) and following the manufacturer’s protocol. The consequent red fluorescent cells were renamed with a suffix “R” (A431-R, MDA-MB-231-R, MDA-MB-453-R, SK-BR-3-R, and HT29-R).

A431-R, MDA-MD-231-R, RAW 264.7, and J558L cells were cultured in Dulbecco’s Modified Eagle’s Medium (DMEM) with 4.5 g/L glucose, L-glutamine, and supplemented with 10% fetal bovine serum (FBS), 100 U/mL penicillin, and 100 μg/mL streptomycin at 37 ºC with 5.0% CO_2_. MDA-MB-453-R and Raji cells were cultured in Roswell Park Memorial Institute (RPMI) medium with L-glutamine and supplemented with 10% FBS, 100 U/mL penicillin, and 100 μg/mL streptomycin at 37 ºC with 5.0% CO_2_. SK-BR-3-R and HT29-R cells were cultured in McCoy’s 5A (Modified) Medium and supplemented with 10% FBS, 100 U/mL penicillin, and 100 μg/mL streptomycin at 37 ºC with 5.0% CO_2_. NK-92 cells were cultured in Minimum Essential Medium α and supplemented with 10% FBS, 100 U/mL penicillin, 1000 U/mL IL-2, and 100 μg/mL streptomycin at 37 ºC with 5.0% CO_2_. HUVEC cells were cultured using the EGM-2™ Endothelial Cell Growth Medium-2 BulletKit™ (Lonza, Cat: CC-3162).

Peripheral blood mononuclear cells (PBMCs) were purified from buffy coats of healthy donors blood samples using Ficoll-Hypaque density gradient centrifugation as previously described^22^ and cultured in ImmunoCult™-XF T Cell Expansion Medium supplemented with 30 U/ml IL-2, 100 U/mL penicillin, and 100 μg/mL streptomycin at 37 ºC with 5.0% CO_2_. The healthy donor blood samples (Donor 9) were purchased from Memorial Blood Centers, Saint Paul, MN. The CD8^+^ T cells were isolated using the Dynabeads™ CD8 Positive Isolation Kit (Invitrogen, Cat: 11333D) and cultured in ImmunoCult™-XF T Cell Expansion Medium supplemented with 30 U/ml IL-2, 100 U/mL penicillin, and 100 μg/mL streptomycin at 37 ºC with 5.0% CO_2_.

### Quantification of cell surface receptors by flow cytometry

The Bangs beads (Bangs Laboratories, Cat: 815A) were collected and resuspended in PBS, and then stained with the fluorescently labeled antibody that detects the corresponding cell surface receptor (anti-EGFR-BV421, Biolegend, Cat: 352911; anti-HER2-BV421, Biolegend, Cat: 324420; anti-EpCAM-AF647, Biolegend, Cat: 118211) at 4 ºC for >30 min, followed by wash steps. 100,000 of the cancer cells were collected and stained with the corresponding antibody at 4 ºC for >30 min, followed by wash steps. The cells and beads were analyzed by flow cytometry. The calibration curve was generated based on the MFI of the beads and the corresponding antibody binding capacity of the beads. The receptor number was calculated based on the MFI of the cells stained with the antibody and the previously generated calibration curve. The confirmation of the CD133 expression on HT29-R cells was carried out with a similar flow cytometry experiment, where the anti-CD133-PE antibody (Biolegend, Cat: 372803) was used to stain the HT29-R cells at 4 ºC for >30 min, followed by wash steps and flow cytometry analysis.

### Expression plasmids and oligonucleotides

gBlock Gene Fragments coding for the αEGFR-Fn3-DHFR^2^-CVIA, αHER2-afb-DHFR^2^-CVIA, αEpCAM-Fn3-DHFR^2^-CVIA, and αCD133-scFv-DHFR^2^ fusion proteins were ordered from Integrated DNA Technologies (IDT) and cloned into the Novagen pET28a(+) vector (EMD Millipore, Cat: 69864-3) via NcoI and XhoI restriction sites. The gene fragment for the DHFR^2^-CVIA protein was generated via site-directed mutagenesis of the gene of αEpCAM-Fn3-DHFR^2^-CVIA protein using a New England Biolabs Q5 Site-Directed Mutagenesis Kit (Cat: E0554S). The ssDNAs containing DBCO functional group were ordered from Integrated DNA Technologies (IDT). The sequences of the protein constructs and ssDNAs are listed in Supplementary Note 1.

### Protein expression and purification

The αEGFR-Fn3-DHFR^2^-CVIA, αHER2-afb-DHFR^2^-CVIA, αEpCAM-Fn3-DHFR^2^-CVIA, αCD133-scFv-DHFR^2^, and DHFR^2^-CVIA fusion proteins were produced in T7 Express Competent *E. coli* cells (New England Biolabs) using 0.5 mM IPTG at 37 ºC for 3-6 hours. The αEGFR-Fn3-DHFR^2^-CVIA and αEpCAM-Fn3-DHFR^2^-CVIA fusion proteins were purified from the soluble fractions of the cell lysate via immobilized metal affinity chromatography (IMAC) using the cobalt column (Thermo Fisher Scientific, Cat: 89964) according to previously reported methods^19^; meanwhile, the αHER2-afb-DHFR^2^-CVIA and DHFR^2^-CVIA fusion proteins were purified from the soluble fractions of the cell lysate via methotrexate affinity chromatography and DEAE ion-exchange chromatography, also according to the previously reported methods^19^. The αCD133-scFv-DHFR^2^ protein was purified from the insoluble fractions of the cell lysate via previously reported denaturation and refolding procedures, followed by Q Sepharose Fast Flow anion exchange chromatography and SEC^22^. Purified protein was analyzed by gel electrophoresis using NuPAGE Bis-Tris protein gels (Thermo Fisher Scientific, Cat: NP0321PK2). DTT (5mM) was added to the samples for gel electrophoresis. Yeast farnesyl transferase (yFTase) was expressed and purified following the previously reported procedures^19,29^.

To prepare the fluorescein-labeled proteins, the protein of interest (10 μM) was incubated with a 10-fold molar excess of NHS-Fluorescein (Thermo Fisher Scientific, Cat: 46410) in PBS for 12 hours at room temperature and then purified through buffer exchange with the Amicon Ultra-0.5 centrifugal filters (10 kDa cutoff, Millipore). The NHS-Fluorescein labeling of the proteins was confirmed by gel electrophoresis, followed by in-gel fluorescent scanning using the Typhoon FLA 9500 (GE Healthcare) and then the Coomassie brilliant blue staining. Gel images were processed in ImageJ.

### Preparation of farnesylated proteins and protein conjugates

Farnesylation reactions were conducted following the previously reported methods. Specifically, a reaction cocktail (typically 500 uL) was prepared with the Dulbecco’s phosphate-buffered saline (DPBS) buffer (Gibco, Cat: 14040141) containing MgCl_2_ (0.5 mM), ZnCl_2_ (10 μM), DTT (5 mM), and the protein of interest (2.5 μM). The mixture was incubated on ice for 0.5 h and the reaction was initiated by the addition of FPP (7.5 μM) or C10-N_3_-OPP (10 μM) with yFTase (200-400 nM) and allowed to proceed for 3-6 h in a 32 ºC water bath. The prenylated protein was subsequently purified by buffer exchange (PBS) with an Amicon Ultra-0.5 centrifugal filter (10 kDa cutoff, Millipore) for cell surface modification or the following conjugation reactions.

For the preparation of protein-drug conjugates, the purified azide containing farnesylated proteins were incubated with a 10-fold molar excess of the DBCO-PEG_4_-VC-PAB-MMAE (ACES Pharma.) at room temperature in the dark for 12 hours, followed by dialysis purification in PBS. Similarly, to prepare the protein-ssDNA conjugates, the azide containing farnesylated proteins were incubated with a 10-fold molar excess of the DBCO-functionalized ssDNA at room temperature in the dark for 12 hours, followed by buffer exchange (PBS) with the Amicon Ultra-0.5 centrifugal filters (10 kDa cutoff, Millipore).

The proteins and protein conjugates were washed into ultrapure water with the Amicon Ultra-0.5 centrifugal filters (10 kDa cutoff, Millipore). 50 μL of each protein (5 μM) or the protein conjugates was characterized by LC-MS using an Orbitrap Elite Hybrid Mass Spectrometer. The data were further processed by the Thermo Scientific™ Protein Deconvolution software.

### CSAN formation and characterization

CSANs were formed by the addition of a 1.1-1.5-fold molar excess of the dimerizer, bisMTX, to a solution of the DHFR^2^ fusion protein monomers (1-2 mL, unless specified otherwise). The oligomerization occurs within minutes after adding bisMTX. CSAN formation was characterized by dynamic light scattering and Cyro-TEM imaging. The hydrodynamic diameters of CSANs were measured by dynamic light scattering with an Anton Paar particle size analyzer (Litesizer 500) and presented as mean value ± standard deviation of at least three measurements. The CSAN samples for cryo-TEM were prepared at 1 μM concentrations in PBS buffer. The CSAN solutions (2.5 μL) were applied to a lacey Formvar/carbon grid (Ted Pella, Inc.; Cat: 01883) in the humidified chamber of a Vitrobot Mark IV (FEI), blotted for 13 seconds, and plunged into liquid ethane for vitrification. Grids were imaged on a Tecnai Spirit G2 BioTWIN (FEI) equipped with an Eagle 2k CCD camera (FEI) under a high tension of 120 kV.

### Cell surface labeling study with CSANs by flow cytometry

The binding specificity of the CSANs to the corresponding cellular receptors was studied by flow cytometry. A431-R cells were chosen as the EGFR^+^ cell line; SKBR-3-R cells were chosen as the HER2^+^ cell line; HT-29-R cells were chosen as the EpCAM^+^ cell line; HT29-R cells were chosen as the CD133^+^ cell line. The cells were harvested and washed with DPBS buffer (Gibco, Cat: 14190144), and aliquots of 10^5^ cells were then resuspended in 100 μL of DPBS solutions containing 0.5 μM of fluorescein-labeled CSANs and incubated for 1 h at 4 ºC. The cells were then pelleted, washed, and resuspended in 0.5 mL of cold DPBS and analyzed using an LSR II flow cytometer (BD Biosciences) at the University Flow Cytometry Resource (UFCR).

For the cell surface modification with farnesylated CSANs, the sender cells were collected from cell culture, pelleted at 350 g for 5 min, and washed with 1 mL DPBS. Aliquots of 10^5^ cells were then incubated in 100 μL of DPBS solutions containing the desired concentrations of fluorescein-labeled farnesylated CSANs for at least 1 h at room temperature with rotation and washed twice with 1 mL cold DPBS to remove unbound CSANs. The modified cells were then resuspended in 0.5 mL of cold DPBS and analyzed using an LSR II flow cytometer (BD Biosciences) at the University Flow Cytometry Resource (UFCR).

### Imaging of surface-bound farnesylated CSANs by fluorescent microscopy

10^5^ Raji cells were harvested and washed with DPBS, followed by modification with the fluorescein-labeled αEGFR-Fn3-Far CSANs (1 μM) for at least 1 h at room temperature with rotation and washed twice with 1 mL cold DPBS to remove unbound CSANs. The cells were then transferred to the 35 mm glass coverslip bottom dish (ibidi, cat: 81158) and imaged by the Nikon Ti-E microscope with an ibidi Stage Top Incubation System.

### Flow cytometry study of non-specific intercellular transfer of farnesylated CSANs

The αEGFR-Fn3-DHFR^2^-CVIA protein was non-specifically labeled with DyLight™ 650 NHS Ester (Thermo Scientific, Cat: 62266) in PBS according to the manufacturer’s protocol. The fluorescently labeled αEGFR-Fn3-DHFR^2^-CVIA (2 μM) was farnesylated following previously described methods, and the αEGFR-Fn3-DHFR^2^-Far protein was oligomerized to form CSANs. 4 × 10^5^ Raji cells were collected and stained with CFSE (Thermo Fisher Scientific, Cat: C34554) according to the manufacturer’s protocol. Meanwhile, 6 × 10^5^ Raji cells were collected and modified with the DyLight™ 650-labeled αEGFR-Fn3-Far CSANs at room temperature with rotation for 1 hour, followed by wash steps with DPBS. The CSAN-modified Raji cells were mixed with the CFSE-stained Raji cells at a 6:4 ratio and were divided into 9 aliquots; each aliquot was incubated in 1 mL of the culture medium. 3 aliquots were analyzed by flow cytometry, and the other 6 aliquots of the cells were returned into the cell culture in a 6-well plate for 24-48 h in the incubator. At 24 h intervals, 3 aliquots of the cells were taken out for flow cytometry analysis. The media was refreshed every 24 h. An LSR II flow cytometer was used for the flow cytometry analysis, and the CSAN^+^/CFSE^+^ (DyLight 650^+^/CFSE^+^) population was quantified as the indicator of non-specific intercellular transfer of the E1-DHFR^2^-Far CSANs.

### Flow cytometry study of the cell-cell CSAN transfer between Raji cells and A431-R cells

Raji cells (6 × 10^4^ per sample) were collected and modified with the fluorescein-labeled αEGFR-Fn3-Far CSANs or fluorescein-labeled DHFR^2^-Far CSANs (1 μM) respectively at room temperature with rotation for 1 hour, followed by wash steps with DPBS. The CSAN-modified Raji cells were mixed with the A431-R cells at a 6:4 ratio and co-cultured in Eppendorf tubes with rotation at 37 ºC (if not otherwise specified). The cells are then analyzed by an LSR II flow cytometer and the CSAN^+^/mKate^+^ (FITC^+^/mKate^+^) cell population was quantified as the indicator of intercellular transfer of CSANs from Raji cells to A431-R cells. Unless otherwise stated, experiments were conducted in triplicate and data are presented as the mean ± standard deviation of three independent trials.

For the transwell assay of the cell-cell CSAN transfer study, Raji cells were collected and modified with the fluorescein-labeled αEGFR-Fn3-Far CSANs (1 μM), followed by wash steps with DPBS. The A431-R cells were collected, and one group of the cells were co-cultured together with the CSAN-modified Raji cells at a 1:1 ratio, while for the other group, the transwell inserts were used to separate the A431-R cells and the CSAN-modified cells. The cells were co-cultured at 37 °C for 30 min. Then the cells were analyzed by flow cytometry. The A431-R cells were gated out and the fluorescence of the fluorescein-labeled CSANs were quantified. All experiments were conducted in triplicate, and data are presented as the mean ± standard deviation of three independent trials.

For the competition binding assay of the cell-cell CSAN transfer study, the Raji cells were collected and modified with the fluorescein-labeled αEGFR-Fn3-Far CSANs (1 μM), followed by wash steps with DPBS. A431-R cells were collected and incubated with different concentrations of unfarnesylated αEGFR-Fn3 CSANs (0-5000 nM) for 10 min, and then were co-cultured in Eppendorf tubes with the CSAN-modified Raji cells at a 6:4 ratio with rotation at 37 °C for 30 min, followed by flow cytometry analysis. All experiments were conducted in triplicate and data are presented as the mean ± standard deviation of three independent trials.

To study the cell-cell CSAN transfer at a low sender-to-receiver ratio, the CSAN-modified Raji cells were co-cultured with A431-R cells at a 1:9 ratio with rotation at 37 °C for 1 h, followed by flow cytometry analysis. In addition, to study the cell-cell CSAN transfer at different temperatures, the CSAN-modified Raji cells were co-cultured with A431-R cells at a 6:4 ratio at 37 °C or 4 °C for 30 min, followed by flow cytometry analysis. All experiments were conducted in triplicates and data are presented as the mean ± standard deviation of three independent trials.

### Flow cytometry study of the CASN-mediated cell-cell interactions during the intercellular CSAN transfer

Raji cells were collected, stained by the Hoechst dye (Invitrogen, Cat: H3570), and modified with the fluorescein-labeled αEGFR-Fn3-Far CSANs (0-2.5 μM) as previously described, followed by wash steps with DPBS. A431-R cells were collected and were co-cultured with the CSAN-modified Raji cells at a 1:1 ratio in tubes with rotation at 37 °C for 1 h, followed by flow cytometry analysis. The Hoechst^+^/mKate^+^ population was quantified to indicate cell-cell interactions, and the FITC^+^/mKate^+^ population was quantified to indicate cell-cell CSAN transfer. All experiments were conducted in triplicates and data are presented as the mean ± standard deviation of three independent trials.

### Kinetics study of the cell-cell CSAN transfer for different CSANs and cell lines

The fluorescein-labeled farnesylated CSANs were prepared following the previously described methods. The Raji cells were collected and modified with the fluorescein-labeled farnesylated CSANs as previously described, followed by wash steps with DPBS. The CSAN-modified Raji cells were co-cultured with the corresponding mKate-expressing receiver cells at a 6:4 ratio with rotation at 37 °C for different times, followed by flow cytometry analysis. All experiments were conducted in triplicates and data are presented as the mean ± standard deviation of three independent trials.

### Recording cell-cell interactions by the C4T approach using flow cytometry

The sender cells were collected and modified with the fluorescein-labeled farnesyl-CSANs (1 μM) at room temperature with rotation for 1 hour, followed by wash steps with DPBS. The receiver cells were collected and co-cultured with the CSAN-modified sender cells in the tubes at a 6:4 ratio with rotation at 37 °C for 30 min, followed by the flow cytometry analysis. The mKate^+^ receiver cells were gated out and the fluorescence of the fluorescein-labeled CSANs was quantified. All experiments were conducted in triplicates and data are presented as the mean ± standard deviation of three independent trials.

### Recording cell-cell interactions by the C4T approach using fluorescent microscopy

1.5 × 10^5^ mKate-expressing target receiver cells were plated in the 35 mm glass coverslip bottom dish (ibidi, cat: 81158) one day prior to the imaging experiment. 10^5^ sender cells were collected and modified with the fluorescein-labeled farnesyl-CSANs at room temperature with rotation for 1 hour, followed by wash steps with DPBS. The CSAN-modified sender cells were added to the cell culture dish of the mKate-expressing cells at 37 °C and then imaged by the Nikon Ti-E microscope with an Ibidi Stage Top Incubation System.

### Flow cytometry and imaging study of the interaction-dependent delivery of MMAE by the C4T approach

To study the interaction-dependent delivery of MMAE by the C4T approach using flow cytometry, the αEGFR-Fn3-DHFR^2^-MMAE was labeled with NHS-Fluorescein and oligomerized with the αEGFR-Fn3-DHFR^2^-Far protein in a 1:3 ratio to form the hybrid αEGFR-Fn3-Far-MMAE CSANs. The Raji cells were collected and modified with the hybrid αEGFR-Fn3-Far-MMAE CSANs (2 μM) at room temperature with rotation for 1 hour, followed by wash steps with DPBS. The CSAN-modified Raji cells were co-cultured with A431-R cells at a 6:4 ratio with rotation at 37 °C or 4 °C for 1 hour, followed by flow cytometry analysis. For the competition binding control, the A431-R cells were pre-incubated with 5 μM of unfarnesylated αEGFR-Fn3 CSANs for 10 min before being co-cultured with the CSAN-modified Raji cells. The cell-cell transfer of the hybrid αEGFR-Fn3-Far-MMAE CSANs from Raji cells to A431-R cells was also imaged by fluorescent microscope following previously described methods used for recording cell-cell interactions.

### Cytotoxicity study of the interaction-dependent delivery of MMAE by the C4T approach

2,500 of A431-R, MDA-MB-231-R, or MDA-MB-453-R cells were plated in the 96-well plates one day before the treatment. The hybrid αEGFR-Fn3-Far-MMAE CSANs were formed by oligomerizing αEGFR-Fn3-DHFR^2^-Far and αEGFR-Fn3-DHFR^2^-MMAE in a 3:1 ratio, while the control hybrid DHFR^2^-MMAE CSANs were formed by oligomerizing DHFR^2^-Far and DHFR^2^-MMAE in a 3:1 ratio. The Raji cells were collected and modified with the CSANs (2 μM) at room temperature with rotation for 1 hour, followed by wash steps with DPBS. The CSAN-modified Raji cells were co-cultured the target cancer cells at a 3:1 ratio in the plate at 37 °C for 2 hours. Then the Raji cells were removed by the medium exchange. The cancer cells were returned to the IncuCyte and cultured for 4 days to quantify cell viability.

### Study of the interaction-dependent delivery of oligonucleotides by the C4T approach

To study the interaction-dependent delivery of ssDNA by the C4T approach using flow cytometry, the αEGFR-Fn3-ssDNA-AF488 protein conjugate was oligomerized with the αEGFR-Fn3-DHFR^2^-Far protein in a 1:3 ratio to form the hybrid αEGFR-Fn3-Far-ssDNA-AF488 CSANs. The Raji cells were collected and modified with the hybrid αEGFR-Fn3-Far-ssDNA-AF488 CSANs (2 μM) at room temperature with rotation for 1 hour, followed by wash steps with DPBS. The CSAN-modified Raji cells were co-cultured with A431-R cells at a 6:4 ratio with rotation at 37 °C or 4 °C for 1 hour, followed by flow cytometry analysis. For the competition binding control, the A431-R cells were pre-incubated with 5 μM of unfarnesylated αEGFR-Fn3 CSANs for 10 min before being co-cultured with the CSAN-modified Raji cells.

To study the specific knockdown of the targeted protein by the C4T approach, several protein-ssDNA conjugates were constructed as previously described, which include the αEGFR-Fn3-DHFR^2^-KDssDNA with an antisense phosphorothioate ssDNA targeting eIF4E, the αEGFR-Fn3-DHFR^2^-CTRLssDNA with a control ssDNA, and the non-targeting DHFR^2^-KDssDNA with the anti-eIF4E phosphorothioate ssDNA. The αEGFR-Fn3-DHFR^2^-KDssDNA was oligomerized with the αEGFR-Fn3-DHFR^2^-Far protein in a 1:3 ratio to form the hybrid αEGFR-Fn3-Far-KDssDNA CSANs. Meanwhile, the non-targeting hybrid DHFR^2^-Far-KDssDNA CSANs were formed by oligomerizing DHFR^2^-KDssDNA and DHFR^2^-Far in a 1:3 ratio, and the hybrid αEGFR-Fn3-Far-CTRLssDNA CSANs were formed by oligomerizing αEGFR-Fn3-DHFR^2^-CTRLssDNA and αEGFR-Fn3-DHFR^2^-Far protein in a 1:3 ratio. 5 × 10^4^ MDA-MB-231 cells were plated in each well of the 24-well plate one day prior to the co-culture experiment. Raji cells were collected and modified with the hybrid CSANs (2 μM) at room temperature with rotation for 1 hour, followed by wash steps with DPBS. The CSAN-modified Raji cells were then co-cultured with MDA-MB-231 cells at a 1:1 ratio at 37 °C for two days and removed from the co-culture through the medium exchange. The MDA-MB-231 cells were then collected, followed by cell lysis with RIPA buffer (Pierce, Cat: 89900) containing complete protease inhibitor cocktail (Roche, Cat:11697498001). The cell lysate samples were collected and normalized to the same concentration, and an equivalent amount of protein lysate (12-15 µg) was electrophoresed on a 4-12% NuPAGE gradient gel and transferred onto low florescent polyvinylidene difluoride membranes (Bio-Rad, Cat: 1704274). Immunoblotting was performed with primary antibodies followed by secondary antibodies with the indicated dilutions: anti-eIF4E, 1:10,000 (R&D Systems, Cat: MAB3228); anti-mouse HRP, 1:1,000 (Invitrogen, Cat: A16072); anti-β-actin, 1:2,000 (Millipore Sigma, Cat: A1978); anti-mouse Alexa Fluor 680, 1;1,000 (Invitrogen, Cat: A32729). When using horseradish peroxidase-conjugated antibodies, West Femto Maximum Sensitivity Substrate (Thermo Scientific, Cat: 34095) was added to the membranes before imaging on an Odyssey Fc Imaging system (Li-Cor). Band intensity was quantified with ImageJ 1.53k.

### Statistical information

Data analysis and data visualization were performed in GraphPad Prism8. Information about error bars, statistical tests, and n values are reported in each figure legend. Unless otherwise stated, experiments were conducted in triplicate, and data are presented as the mean ± standard deviation of three independent trials. In the CSAN transfer kinetics study, the equations of the linear regression lines were generated for the calculation of the T_50_ values (for the study with A431-R cells, the datapoints within the first 10 minutes were used for linear regression analysis), and the slopes of the linear regression lines are statistically different. Differences between means were compared using the unpaired two-tailed Student’s t-tests, and a P-value <0.05 is denoted in graphics with an (*), P < 0.01 is denoted with (**), P < 0.001 is denoted with (***), and P < 0.0001 is denoted with (****).

## Data Availability

The source data for all graphs are provided in the attached Source Data.xlsx file. Raw data files are available from the corresponding author on reasonable request. Source data are provided with this paper.

## Acknowledgements

This work was supported by GM084152 (M.D.D.), CA185627 (C.R.W.), CA247681 (C.R.W), NSF Grant ECCS-2025124 to the Minnesota Nano Center and the Doctoral Dissertation Fellowship from the University of Minnesota. We thank Dr. Robert Hafner for his help with the cryo-TEM experiments, and these experiments were carried out in the Characterization Facility, University of Minnesota, which receives partial support from NSF through the MRSEC program. We thank Dr. Yingchun Zhao for his help with the LC-MS experiments that were conducted in the Masonic Cancer Center’s Analytical Biochemistry Shared Resource. We thank Dr. Mark Sanders for his help with the fluorescent imaging experiments, which were carried out in the University Imaging Center, University of Minnesota. We thank Jian Tang for his help with the western blot experiments. We thank Dr. Bruce Walcheck for his generous offer of the J558L cell line.

## Supplementary Information

