## Supplemental Info for "Macro-Chemical Biology: Engineering Biomimetic Trogocytosis with Farnesylated Chemically Self-Assembled Nanorings"

### A Universal Non-Genetic Method for Engineering Cell-Cell Cargo Transfer with Lipidated Chemically Self-Assembled Nanorings

Yiao Wang[1], Lakmal Rozumalski[2], Caitlin Lichtenfels[2], Jacob Petersberg[2], Ozgun Kilic[2], Mark D. Distefano[1,2]✉ & Carston R. Wagner[1,2]✉

[1]Department of Chemistry, University of Minnesota, Minneapolis, Minnesota, USA

[2]Department of Medicinal Chemistry, University of Minnesota, Minneapolis, Minnesota, USA

University of Minnesota

Department of Medicinal Chemistry

2231 6th Street S.E.

Cancer & Cardiovascular Research Building

Minneapolis, Minnesota 55455, USA

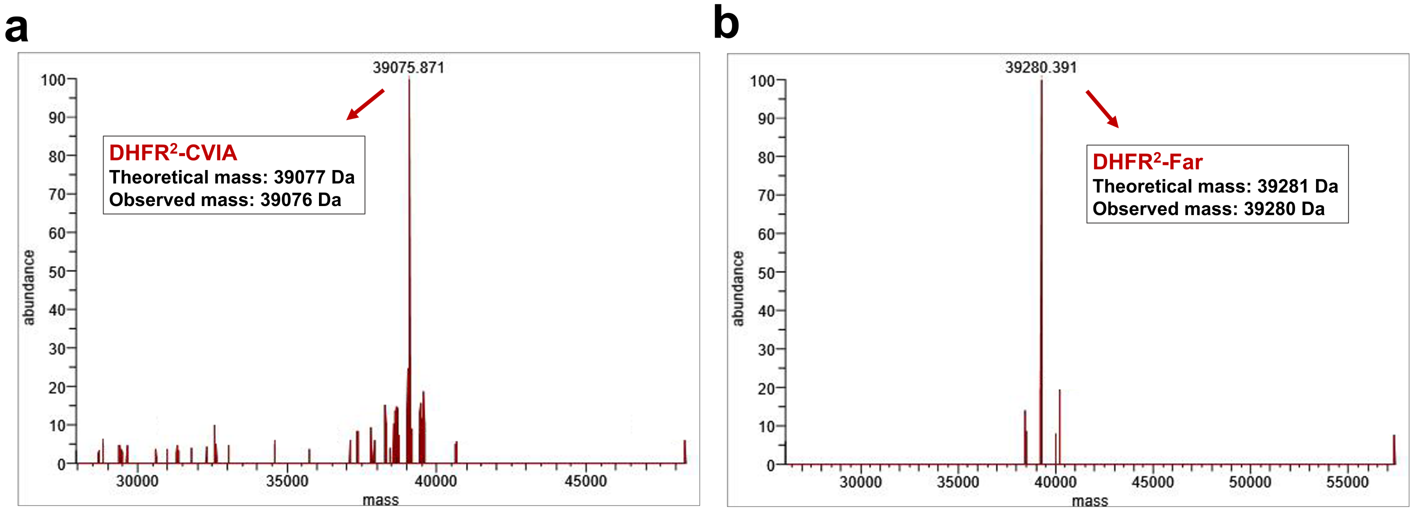

**Supplementary Figure 1. LC-MS spectra of DHFR$^2$-CVIA protein and DHFR$^2$-Far protein.** LC-MS (Orbitrap Elite) was used to characterize the mass of **a** DHFR$^2$-CVIA protein and **b** DHFR$^2$-Far protein, which confirmed the farnesylation of the DHFR$^2$-CVIA protein.

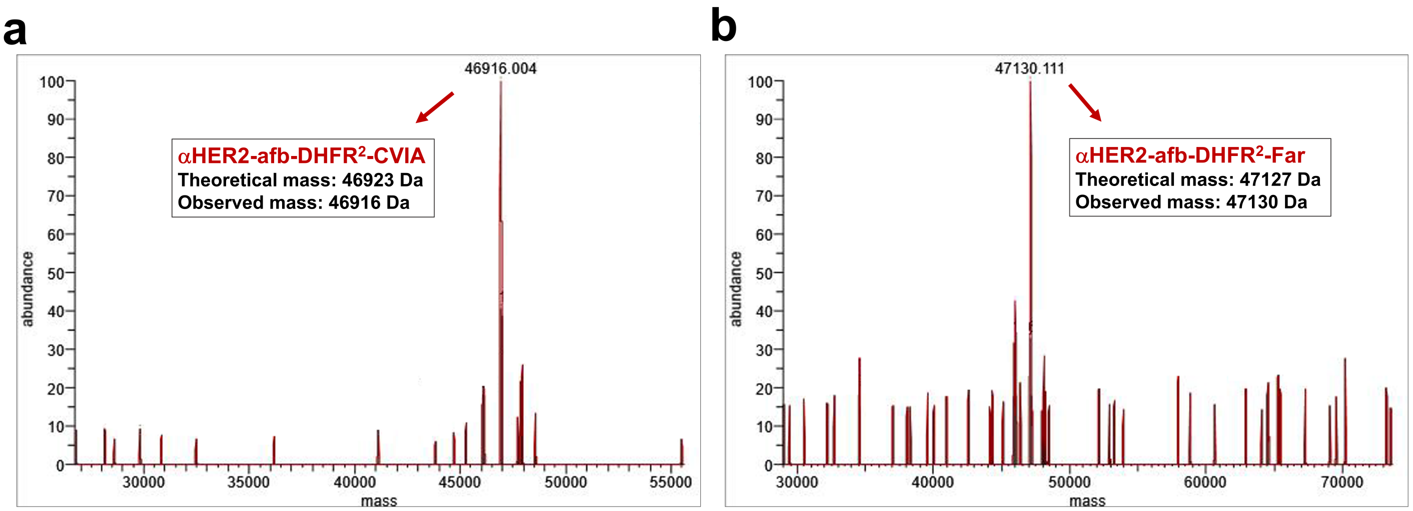

**Supplementary Figure 2. LC-MS spectra of H2-DHFR$^2$-CVIA protein and H2-DHFR$^2$-Far protein.** LC-MS (Orbitrap Elite) was used to characterize the mass of **a** H2-DHFR$^2$-CVIA protein and **b** H2-DHFR$^2$-Far protein, which confirmed the farnesylation of the H2-DHFR$^2$-CVIA protein.

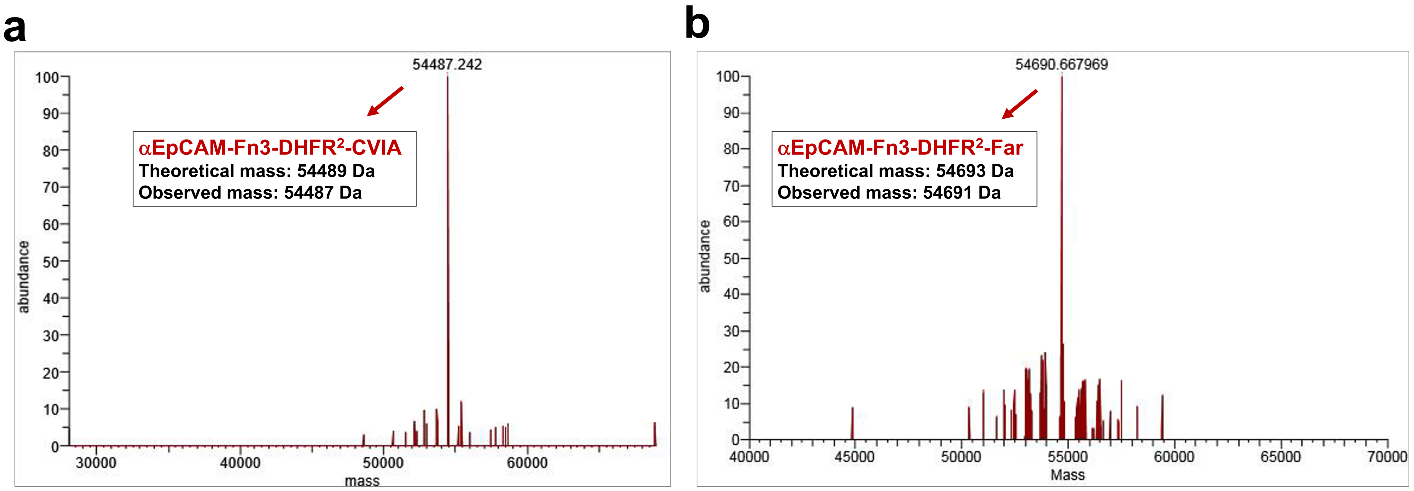

**Supplementary Figure 3. LC-MS spectra of Ep-DHFR$^2$-CVIA protein and Ep-DHFR$^2$-Far protein.** LC-MS (Orbitrap Elite) was used to characterize the mass of **a** Ep-DHFR$^2$-CVIA protein and **b** Ep-DHFR$^2$-Far protein, which confirmed the farnesylation of the Ep-DHFR$^2$-CVIA protein.

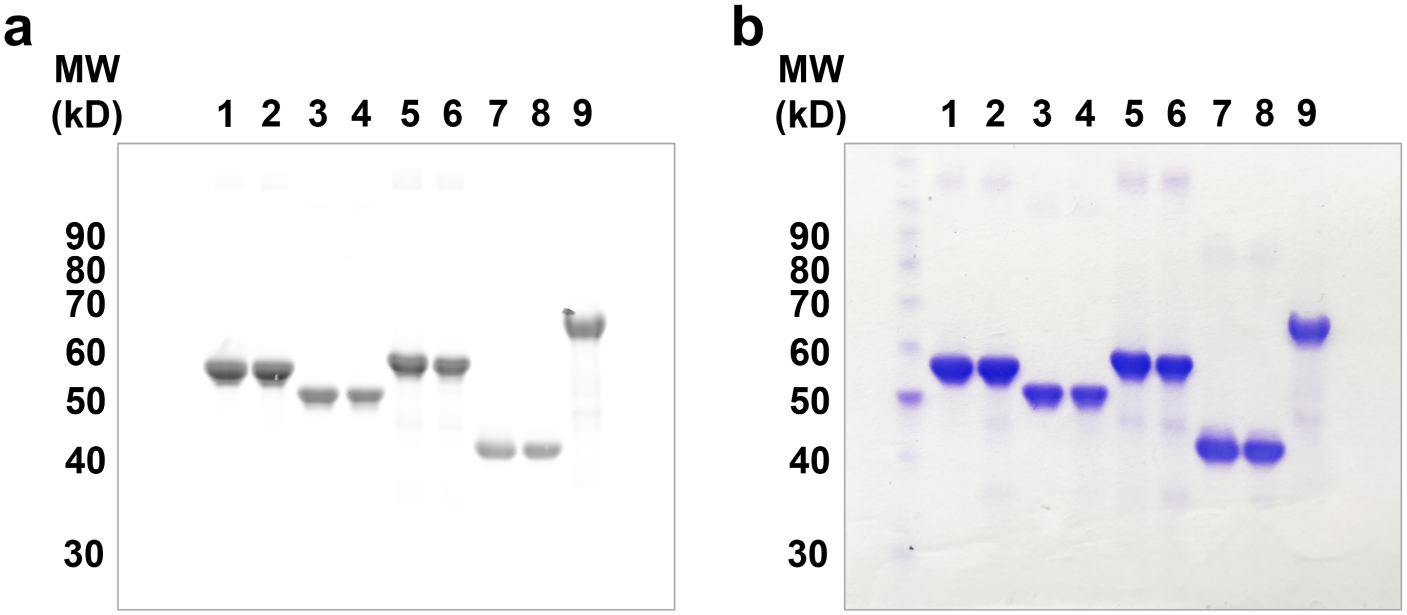

**Supplementary Figure 4. SDS-PAGE gel of the fluorescein-labeled DHFR$^2$ fusion proteins. a** The fluorescein-labeled DHFR$^2$ fusion proteins were characterized by gel electrophoresis, and the fluorescent labeling of the proteins was confirmed by fluorescent scanning of the gel (FITC channel). **b** The SDS-PAGE gel of the fluorescein-labeled DHFR$^2$ fusion proteins was then stained by Coomassie brilliant blue and imaged. Lane 1: fluorescein-labeled E1-DHFR$^2$-CVIA; lane 2: fluorescein-labeled E1-DHFR$^2$-Far; lane 3: fluorescein-labeled H2-DHFR$^2$-CVIA; lane 4: fluorescein-labeled H2-DHFR$^2$-Far; lane 5: fluorescein-labeled Ep-DHFR$^2$-CVIA; lane 6: fluorescein-labeled Ep-DHFR$^2$-Far; lane 7: fluorescein-labeled DHFR$^2$-CVIA; lane 8: fluorescein-labeled DHFR$^2$-Far; lane 9: fluorescein-labeled CD133-DHFR$^2$.

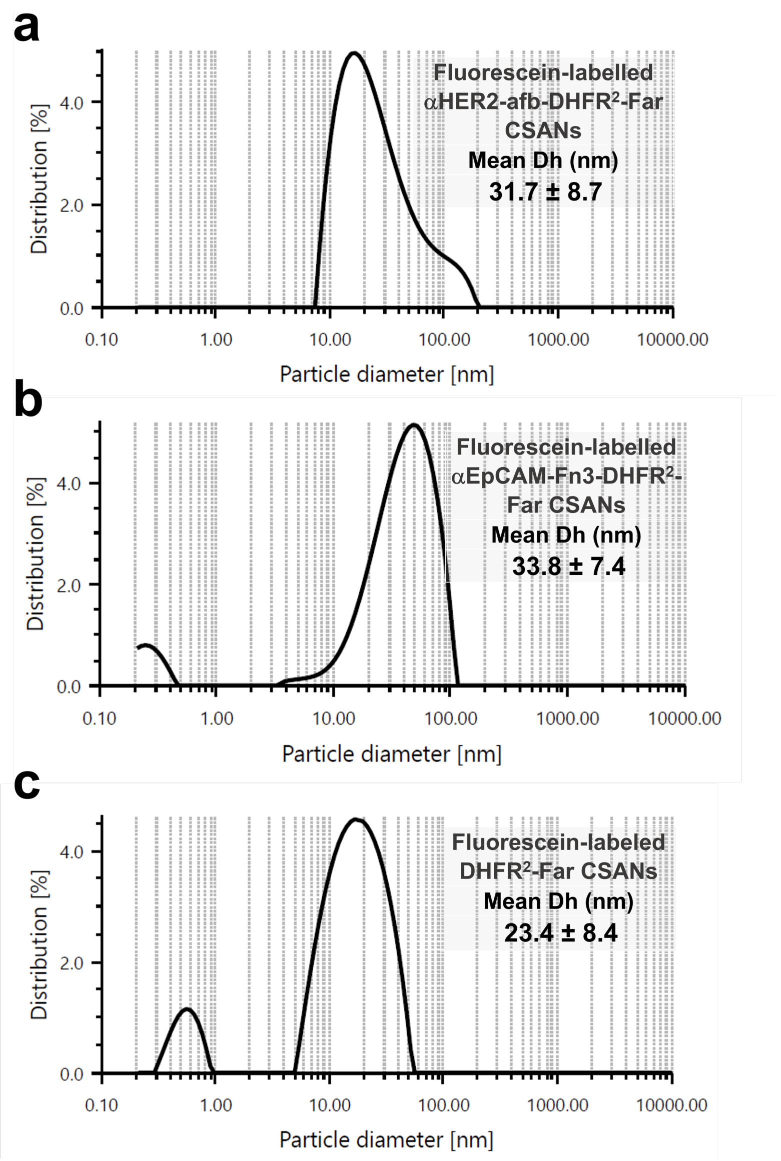

**Supplementary Figure 5. Hydrodynamic diameters of the farnesylated CSANs were measured by dynamic light scattering analysis.** The dynamic light scattering was used to measure the hydrodynamic diameter of **a** fluorescein-labeled H2-DHFR$^2$-Far CSANs (31.7 ± 8.7 nm) **b** fluorescein-labeled Ep-DHFR$^2$-Far CSANs (33.8 ± 7.4 nm) **c** fluorescein-labeled DHFR$^2$-Far CSANs (23.4 ± 8.4 nm).

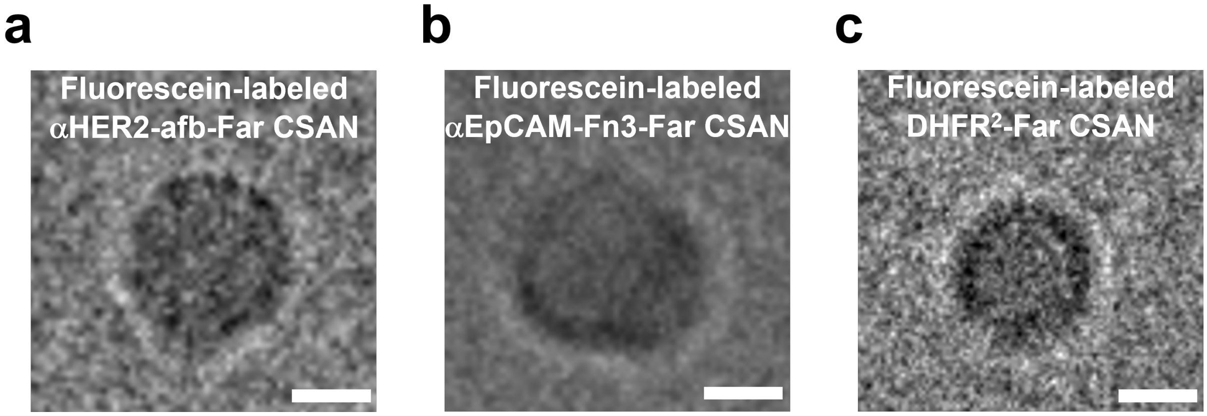

**Supplementary Figure 6. Cryo-TEM imaging of the farnesylated CSANs.** The fluorescein-labeled H2-DHFR$^2$-Far protein, fluorescein-labeled Ep-DHFR$^2$-Far protein, and fluorescein-labeled DHFR$^2$-Far protein was self-assembled into the corresponding nanorings by bisMTX and the CSANs were characterized by cryo-TEM imaging. The morphology of the nanorings exhibited the uniform circular conformation, and the sizes of the constructs are comparable to the DLS data (scale bar, 10 nm).

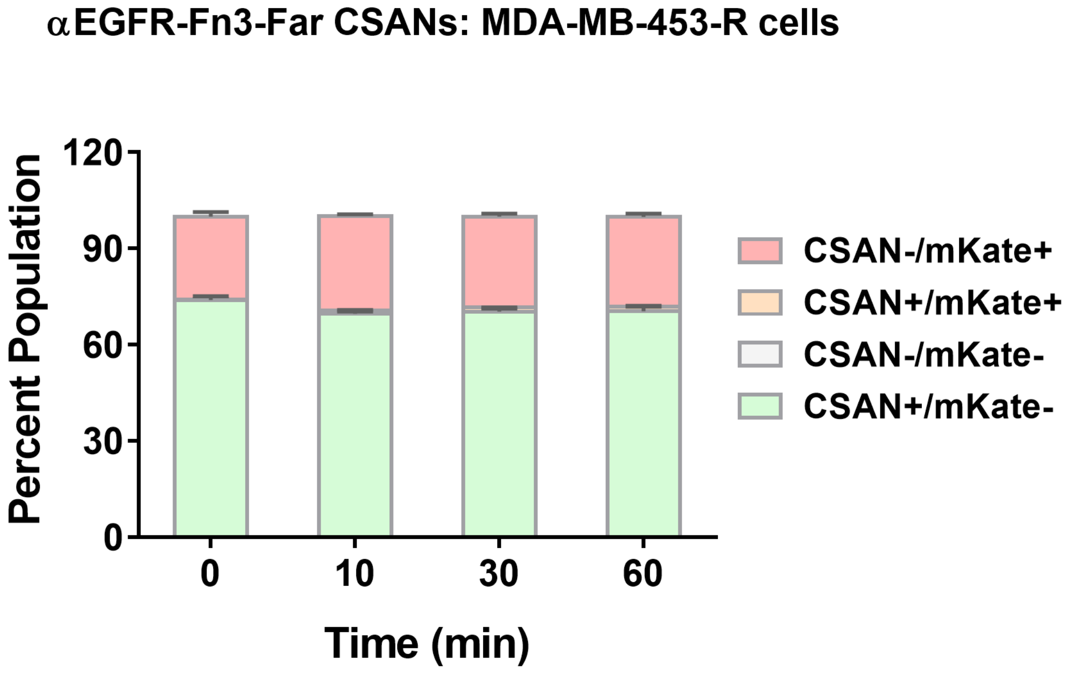

**Supplementary Figure 7. Flow cytometry study of the transfer kinetics of the E1-DHFR$^2$-Far CSANs from Raji cells to MDA-MB-453-R cells.** The CSAN-modified Raji cells were co-cultured with mKate-expressing EGFR-negative MDA-MB-453-R cells at a 6:4 ratio with rotation at 37 °C for 0-60 mins, followed by flow cytometry analysis. No obvious cell-cell CSAN transfer was observed during the co-culture.

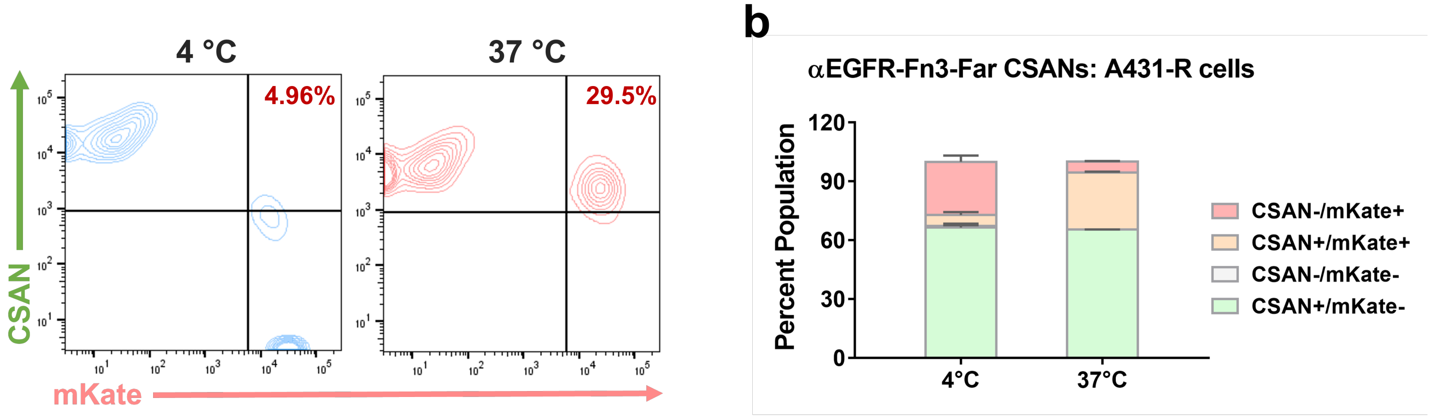

**Supplementary Figure 8. Flow cytometry study of the cell-cell CSAN transfer at different temperatures. a** The representative flow cytometry plots of the cell-cell CSAN transfer study at different temperatures (4 °C vs 37 °C). Raji cells were modified with E1-DHFR$^2$-Far CSANs and co-cultured with A431-R cells at 4 °C or 37 °C for 30 minutes with rotation and the CSAN transfer was then detected and quantified by flow cytometry. The quantitative data of the flow cytometry analysis are presented in **b** the percent cell populations of the cell co-cultures. A significantly higher level of CSAN transfer was observed when cells were co-cultured at 37 °C, indicating the CSAN transfer is associated with the energy-dependent receptor internalization process of the receiver cells. For **b**, data are represented as mean values ± SD (from n=3 independent experimental replicates). In some instances, small error bars are obscured by the symbols denoting the mean value. Source data are provided as a Source Data file.

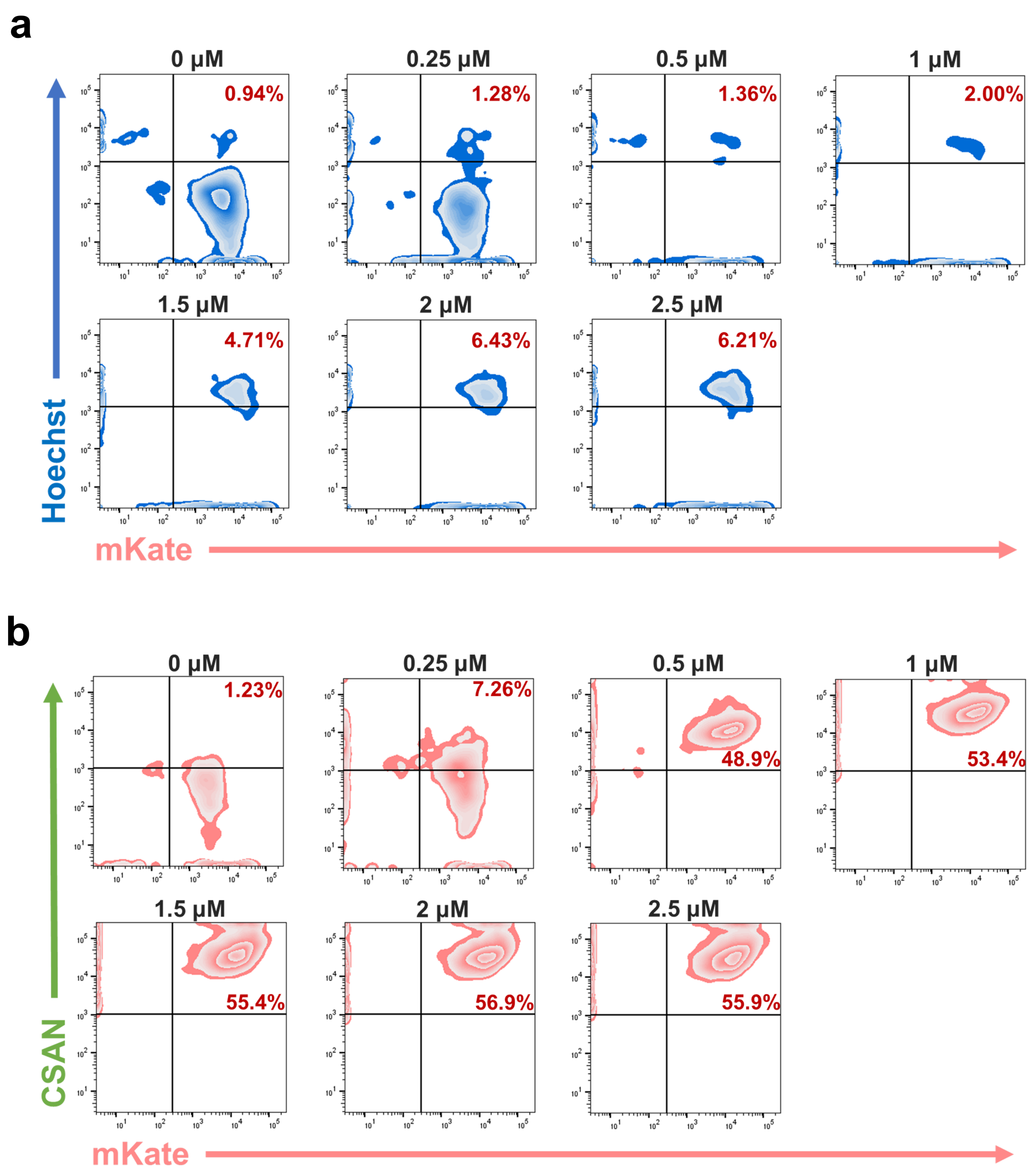

**Supplementary Figure 9. Flow cytometry study showed the CSANs were able to transfer to receiver cells without mediating cell-cell interactions between sender cells and receiver cells. a** Raji cells were stained with Hoechst dye as a marker and modified with different concentrations (0-2.5 μM) of fluorescein-labeled E1-DHFR$^2$-Far CSANs, followed by co-culture with the mKate-expressing A431-R cells. The cells were analyzed by flow cytometry after the co-culture, and the cell-cell interactions were quantified by measuring the percentage of Hoechst$^+$/mKate$^+$ double-positive population, which indicates cell-cell clusters. The CSANs did not induce significant cell-cell interactions at low CSAN concentrations (< 1 μM) for sender cell modifications. **b** The CSAN transfer between sender cells and receiver cells was quantified by measuring the percentage of CSAN$^+$/mKate$^+$ double-positive populations. The CSANs were effectively transferred to the receiver cells even at low CSAN concentrations (< 1 μM) for sender cell modifications.

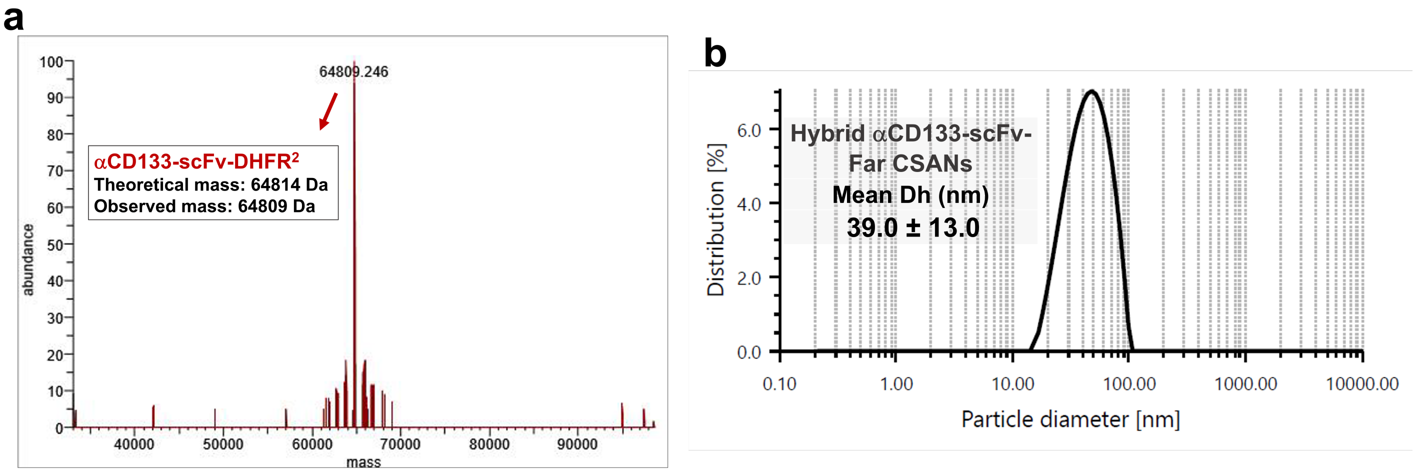

**Supplementary Figure 10. Characterization of CD133-DHFR$^2$ protein and the hybrid DHFR$^2$-Far/CD133-DHFR$^2$ CSANs. a** The CD133-DHFR$^2$ protein was characterized by LC-MS (Orbitrap Elite). **b** The hybrid CD133-DHFR$^2$/DHFR$^2$-Far CSANs were formed by oligomerizing the CD133-DHFR$^2$ protein and the DHFR$^2$-Far protein at a 1:1 ratio. The hydrodynamic diameter of the hybrid CD133-DHFR$^2$/DHFR$^2$-Far CSANs was measured to be 39.0 ± 13.0 nm.

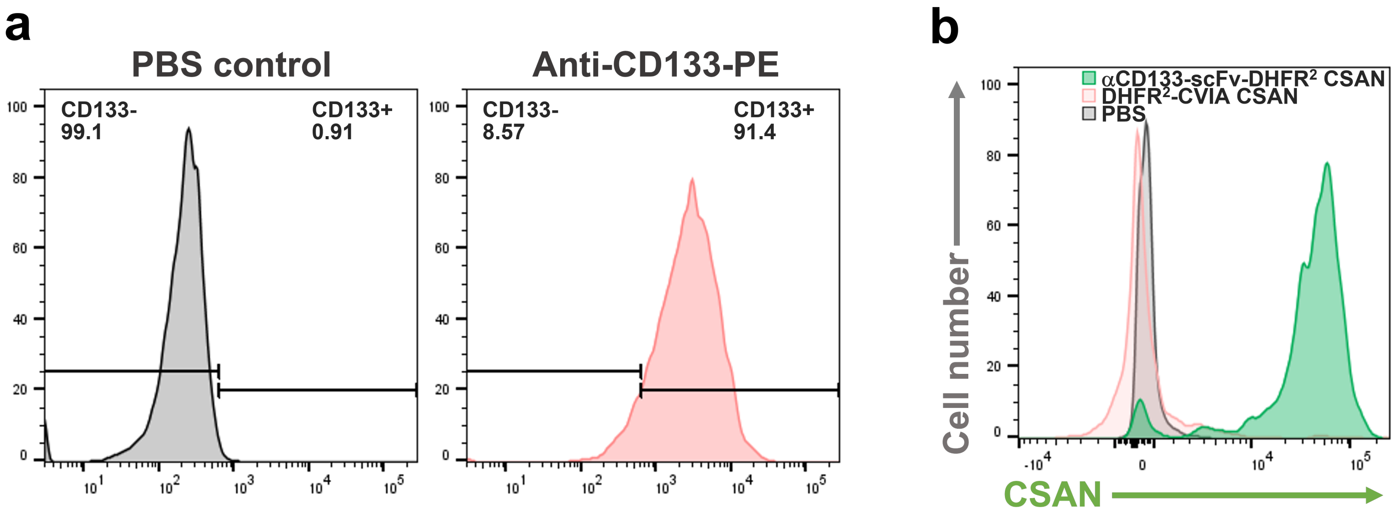

**Supplementary Figure 11. The CD133-DHFR$^2$ CSANs were demonstrated to specifically bind to CD133$^+$ HT29-R cells. a** HT29-R cells were confirmed to be CD133$^+$ by flow cytometry, where the anti-CD133-PE antibody was used to detect the CD133, and over 90% of the HT29-R cells were shown to express CD133. **b** The fluorescein-labeled CD133-DHFR$^2$ CSANs were demonstrated to specifically bind to HT29-R cells by flow cytometry.

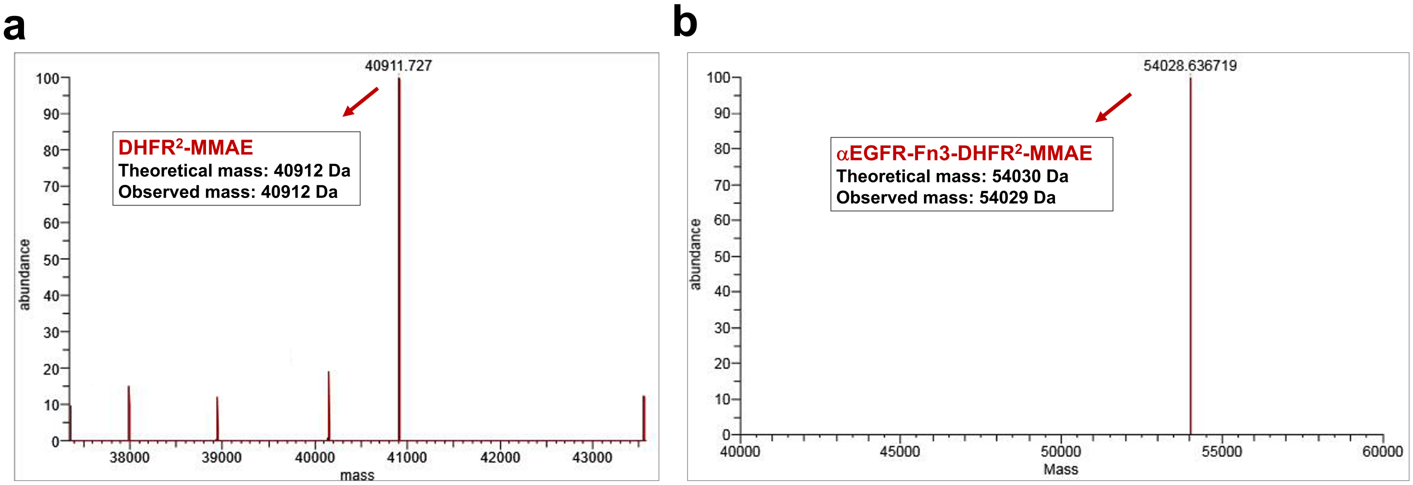

**Supplementary Figure 12. LC-MS spectra of the protein-drug conjugates.** The construction of **a** DHFR$^2$-MMAE protein-drug conjugate and **b** E1-DHFR$^2$-MMAE protein-drug conjugate was confirmed by LC-MS (Orbitrap Elite).

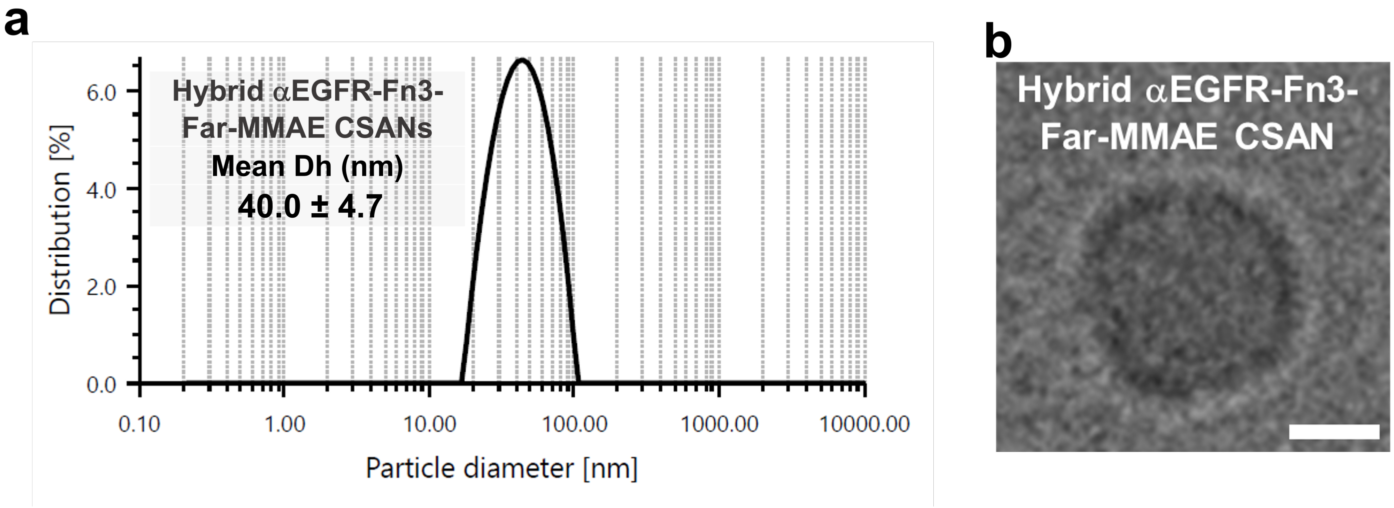

**Supplementary Figure 13. Characterization of the hybrid E1-DHFR$^2$-Far/E1-DHFR$^2$-MMAE CSANs. a** The hybrid E1-DHFR$^2$-Far/E1-DHFR$^2$-MMAE CSANs were formed by oligomerizing the E1-DHFR$^2$-Far protein and the E1-DHFR$^2$-MMAE protein. The hydrodynamic diameter of the hybrid E1-DHFR$^2$-Far/E1-DHFR$^2$-KDssDNA CSANs was measured to be 40.0 ± 4.7 nm. **b** The hybrid E1-DHFR$^2$-Far/E1-DHFR$^2$-MMAE CSANs were characterized by Cryo-TEM imaging. The morphology of the nanorings exhibited a uniform circular conformation, and the size of the constructs is comparable to the DLS data (scale bar, 10 nm).

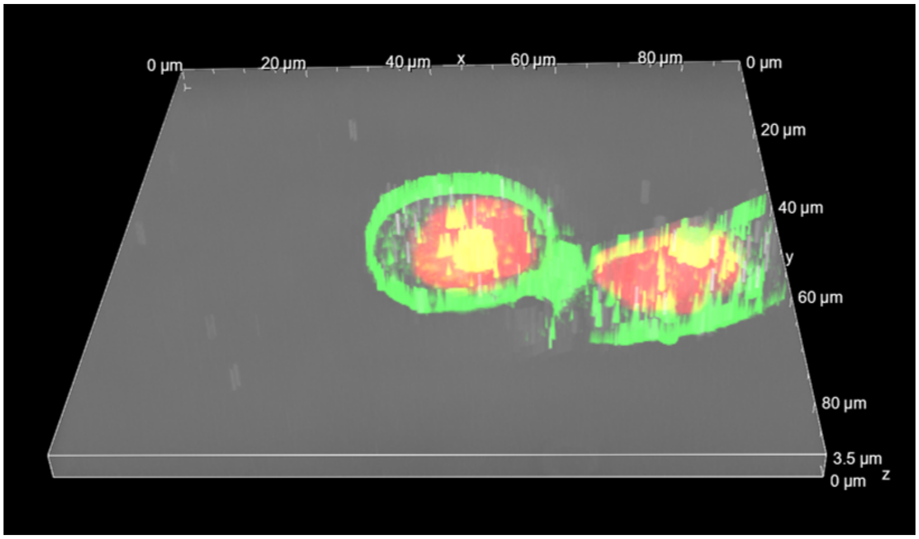

**Supplementary Figure 14. 3-dimentional display of the internalization and localization of the fluorescein-labeled αEGFR-Fn3-DHFR$^2$-MMAE CSANs in A431-R cells by fluorescent microscopy imaging.** The A431-R cells were incubated with 0.5 µM of the fluorescein-labeled aEGFR-Fn3-DHFR$^2$-MMAE CSANs at 37 °C for 1 h and then imaged by an Eclipse Ti-E Wide Field Deconvolution Inverted Microscope (Nikon Instruments, Inc.) using the Z-stack mode. The CSANs are shown in green and the nucleus is shown in red. The green punctate spots indicate the internalized CSANs and the yellow punctate spots indicate the localization of the CSANs in the red nuclei of the A431-R cells.

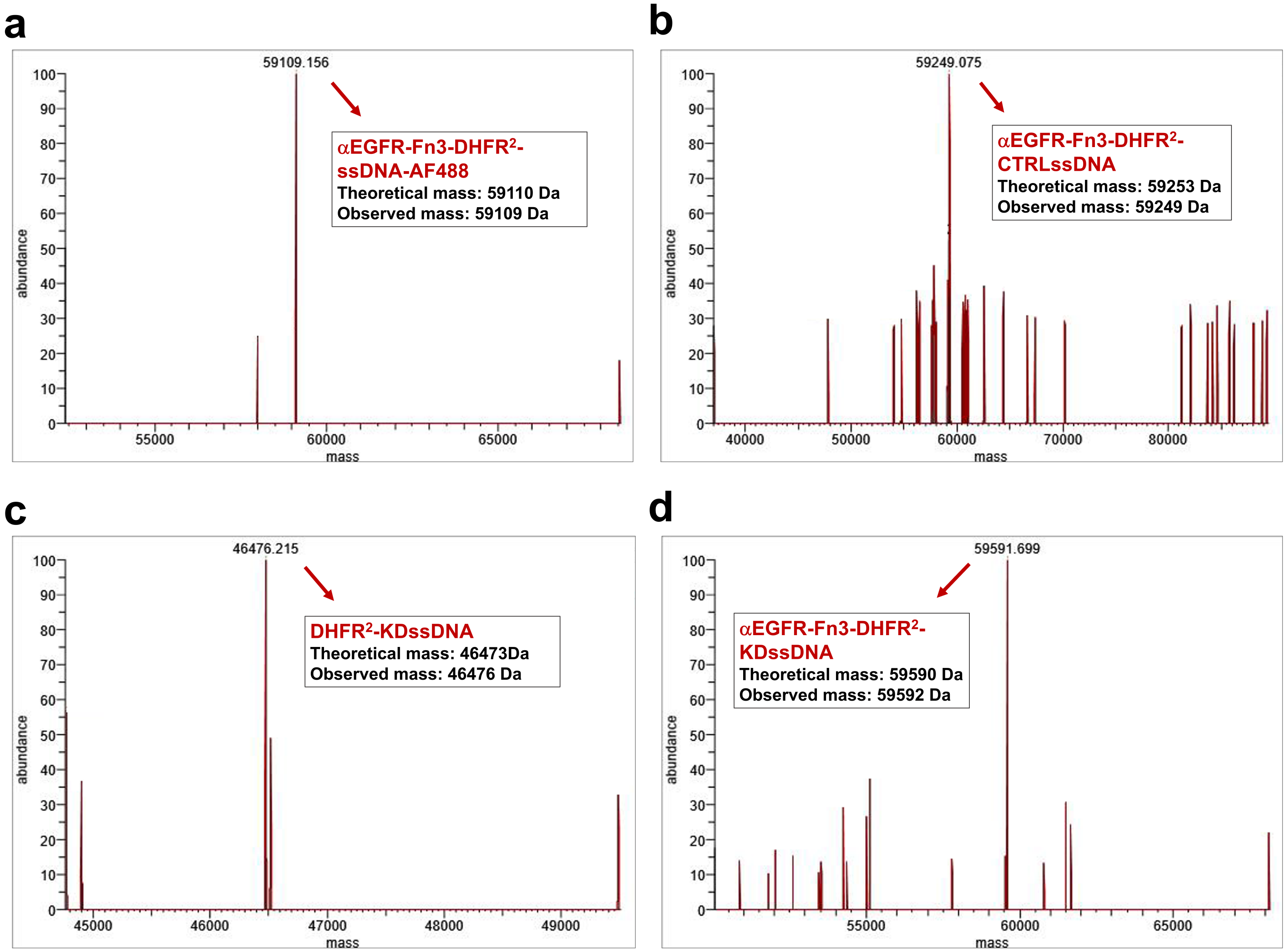

**Supplementary Figure 15. LC-MS spectra of the E1-DHFR²-ssDNA protein-oligonucleotides conjugates. a** The construction of E1-DHFR²-ssDNA-AF488 protein-oligonucleotides conjugate was confirmed by LC-MS (Orbitrap Elite). **b** The construction of E1-DHFR²-CTRLssDNA protein-oligonucleotides conjugate was confirmed by LC-MS (Orbitrap Elite). **c** The construction of DHFR²-KDssDNA protein-oligonucleotides conjugate was confirmed by LC-MS (Orbitrap Elite). **d** The construction of E1-DHFR²-KDssDNA protein-oligonucleotides conjugate was confirmed by LC-MS (Orbitrap Elite).

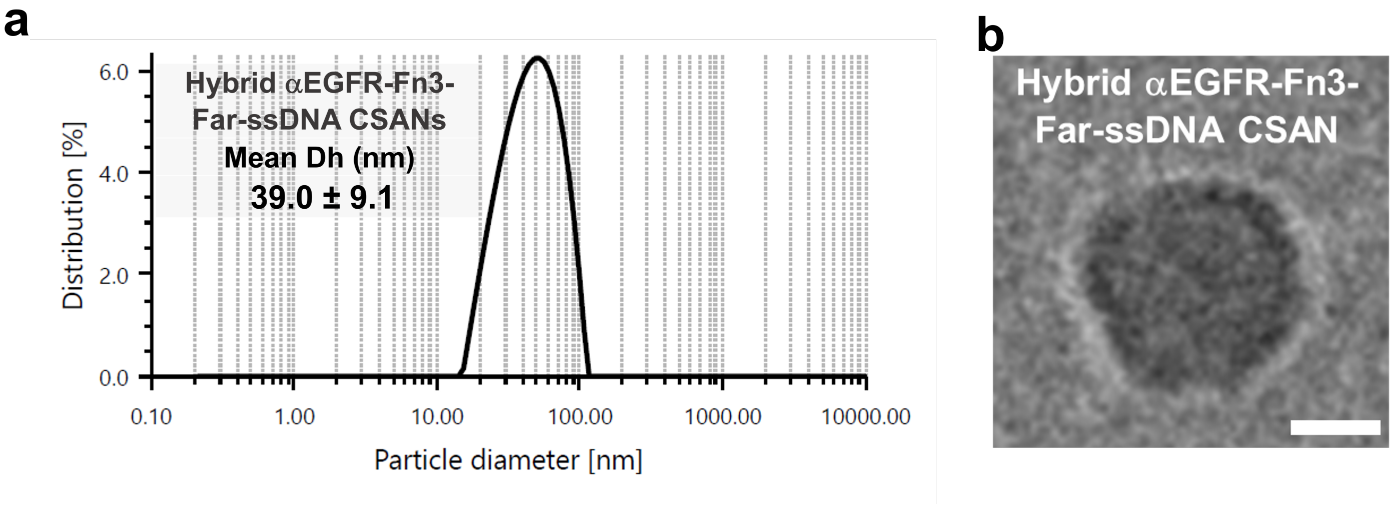

**Supplementary Figure 16. Characterization of the hybrid E1-DHFR$^2$-Far/E1-DHFR$^2$-ssDNA CSANs. a** The hybrid E1-DHFR$^2$-Far/E1-DHFR$^2$-ssDNA-AF488 CSANs were formed by oligomerizing the E1-DHFR$^2$-Far protein and the E1-DHFR$^2$-ssDNA-AF488 protein. The hydrodynamic diameter of the hybrid E1-DHFR$^2$-Far/E1-DHFR$^2$-ssDNA-AF488 CSANs was measured to be 39.0 ± 9.1 nm. **b** The hybrid E1-DHFR$^2$-Far/E1-DHFR$^2$-ssDNA-AF488 CSANs were characterized by Cryo-TEM imaging. The morphology of the nanorings exhibited a uniform circular conformation, and the size of the constructs is comparable to the DLS data (scale bar, 10 nm).

**Supplementary Note 1. Gene sequence of DHFR$^2$ fusion protein constructs and ssDNAs.**

***DHFR$^2$-CVIA*：**

GGCATCAGTCTGATTGCGGCGTTAGCGGTAGATCGCGTTATCGGCATGGAAAACGCCATGCCGTGGAACCTGCCTGCC
GATCTCGCCTGGTTTAAACGCAACACCTTAAATAAACCCGTGATTATGGGCCGCCATACCTGGGAATCAATCGGTCGTC
CGTTGCCAGGACGCAAAAATATTATCCTCAGCAGTCAACCGGGTACGGACGATCGCGTAACGTGGGTGAAGTCGGTGG
ATGAAGCCATCGCGGCGGCTGGTGACGTACCAGAAATCATGGTGATTGGCGGCGGTCGCGTTTATGAACAGTTCTTGC
CAAAAGCGCAAAAACTGTATCTGACGCATATCGACGCAGAAGTGGAAGGCGACACCCATTTCCCGGATTACGAGCCG
GATGACTGGGAATCGGTATTCAGTGAATTCCACGATGCTGATGCGCAGAACTCTCACAGCTATAGCTTTGAGATTCTGG
AGCGGCGGGGCGGCATTAGCCTTATTGCCGCCTTAGCGGTTGATCGCGTGATCGGAATGGAGAACGCAATGCCCTGGA
ATCTTCCGGCAGACCTTGCCTGGTTCAAACGCAACACTTTAAACAAGCCTGTCATTATGGGCCGTCACACATGGGAGTC
AATTGGTCGTCCCCTGCCTGGGCGCAAAAATATCATCTTGTCCTCGCAGCCTGGGACAGATGATCGCGTTACATGGGTG
AAGTCCGTAGACGAAGCGATTGCCGCTGCCGGCGATGTGCCCGAGATTATGGTAATCGGGGGAGGGCGTGTTTACGAA
CAATTTCTGCCCAAAGCTCAGAAATTATACCTGACGCACATCGACGCGGAGGTCGAAGGTGACACACACTTTCCAGAT
TATGAGCCTGATGATTGGGAATCCGTTTTCTCAGAATTTCATGACGCGGATGCTCAAAACTCGCACTCGTACTCTTTTG
AAATTTTAGAGCGCCGTGGCGGATCTGGAGGAAGTGGCGGTGACTACAAAGACGACGATGATAAGGGCGGCTCAGGT
GGTTCCGGTGGCAAAAAGAAAAAGAAAAAGACCTGTGTCATCGCCTAGTGA

***αEGFR-Fn3-DHFR$^2$-CVIA:***

ATGGACTACAAAGACGACGATGATAAGGGCGGATCTGGAGGAAGTGGCGGTATGGGTGTCTCTGACGTCCCGCGTGA
CCTGGAGGTTGTTGCAGCGACCCCAACTAGCCTTCTTATCAGCTGGGATAGCGGTCGTGGTTCTTATCAATACTACCGG
ATCACTTACGGAGAAACAGGAGGAAATAGCCCTGTTCAGGAGTTCACTGTGCCTGGTCCTGTACACACTGCTACCATC
AGCGGCCTTAAACCTGGAGTAGATTATACCATCACTGTGTATGCTGTCACTGACCATAAGCCTCATGCTGATGGACCTC
ACACTTATCATGAATCTCCAATTTCCATCAATTACCGTACAGATATTGATCGTCCTTCTCAAGGTGGTAGTGGCATCAG
TCTGATTGCGGCGTTAGCGGTAGATCGCGTTATCGGCATGGAAAACGCCATGCCGTGGAACCTGCCTGCCGATCTCGC
CTGGTTTAAACGCAACACCTTAAATAAACCCGTGATTATGGGCCGCCATACCTGGGAATCAATCGGTCGTCCGTTGCCA
GGACGCAAAAATATTATCCTCAGCAGTCAACCGGGTACGGACGATCGCGTAACGTGGGTGAAGTCGGTGGATGAAGC
CATCGCGGCGGCTGGTGACGTACCAGAAATCATGGTGATTGGCGGCGGTCGCGTTTATGAACAGTTCTTGCCAAAAGC
GCAAAAACTGTATCTGACGCATATCGACGCAGAAGTGGAAGGCGACACCCATTTCCCGGATTACGAGCCGGATGACTG
GGAATCGGTATTCAGTGAATTCCACGATGCTGATGCGCAGAACTCTCACAGCTATAGCTTTGAGATTCTGGAGCGGCG
GGGCGGCATTAGCCTTATTGCCGCCTTAGCGGTTGATCGCGTGATCGGAATGGAGAACGCAATGCCCTGGAATCTTCC
GGCAGACCTTGCCTGGTTCAAACGCAACACTTTAAACAAGCCTGTCATTATGGGCCGTCACACATGGGAGTCAATTGG
TCGTCCCCTGCCTGGGCGCAAAAATATCATCTTGTCCTCGCAGCCTGGGACAGATGATCGCGTTACATGGGTGAAGTCC
GTAGACGAAGCGATTGCCGCTGCCGGCGATGTGCCCGAGATTATGGTAATCGGGGGAGGGCGTGTTTACGAACAATTT
CTGCCCAAAGCTCAGAAATTATACCTGACGCACATCGACGCGGAGGTCGAAGGTGACACACACTTTCCAGATTATGAG
CCTGATGATTGGGAATCCGTTTTCTCAGAATTTCATGACGCGGATGCTCAAAACTCGCACTCGTACTCTTTTGAAATTTT
AGAGCGCCGTGGCGGCTCAGGTGGTTCCGGTGGCCATCATCATCATCATCACGGCGGCTCAAAAAAGAAAAAGAAAA
AGACCTGTGTCATCGCCTAGTGA

##### *αHER2-afb-DHFR$^2$-CVIA:*

ATGGACTACAAAGACGACGATGATAAGGGCGGATCTGGAGGAAGTGGCGGTATGGCGGAGGCGAAGTACGCGAAAG
AAATGCGTAACGCGTATTGGGAGATCGCGCTGCTGCCGAACCTGACCAACCAGCAAAAGCGTGCGTTCATTCGTAAAC
TGTACGACGATCCGAGCCAGAGCAGCGAGCTGCTGAGCGAAGCGAAGAAACTGAACGACAGCCAAGCGCCGAAGGG
CGGCGGTAGTGGCGGTGGCAGCGGTGGCGGTAGCGGCGGTGGCATCAGTCTGATTGCGGCGTTAGCGGTAGATCGCGT
TATCGGCATGGAAAACGCCATGCCGTGGAACCTGCCTGCCGATCTCGCCTGGTTTAAACGCAACACCTTAAATAAACC
CGTGATTATGGGCCGCCATACCTGGGAATCAATCGGTCGTCCGTTGCCAGGACGCAAAAATATTATCCTCAGCAGTCA
ACCGGGTACGGACGATCGCGTAACGTGGGTGAAGTCGGTGGATGAAGCCATCGCGGCGGCTGGTGACGTACCAGAAA
TCATGGTGATTGGCGGCGGTCGCGTTTATGAACAGTTCTTGCCAAAAGCGCAAAAACTGTATCTGACGCATATCGACG
CAGAAGTGGAAGGCGACACCCATTTCCCGGATTACGAGCCGGATGACTGGGAATCGGTATTCAGTGAATTCCACGATG
CTGATGCGCAGAACTCTCACAGCTATAGCTTTGAGATTCTGGAGCGGCGGGGCGGCATTAGCCTTATTGCCGCCTTAGC
GGTTGATCGCGTGATCGGAATGGAGAACGCAATGCCCTGGAATCTTCCGGCAGACCTTGCCTGGTTCAAACGCAACAC
TTTAAACAAGCCTGTCATTATGGGCCGTCACACATGGGAGTCAATTGGTCGTCCCCTGCCTGGGCGCAAAAATATCATC
TTGTCCTCGCAGCCTGGGACAGATGATCGCGTTACATGGGTGAAGTCCGTAGACGAAGCGATTGCCGCTGCCGGCGAT
GTGCCCGAGATTATGGTAATCGGGGGAGGGCGTGTTTACGAACAATTTCTGCCCAAAGCTCAGAAATTATACCTGACG
CACATCGACGCGGAGGTCGAAGGTGACACACACTTTCCAGATTATGAGCCTGATGATTGGGAATCCGTTTTCTCAGAA
TTTCATGACGCGGATGCTCAAAACTCGCACTCGTACTCTTTTGAAATTTTAGAGCGCCGTGGCGGCTCAGGTGGTTCCG
GTGGCAAAAAGAAAAAGAAAAAGACCTGTGTCATCGCCTAGTGA

##### *αEpCAM-Fn3-DHFR$^2$-CVIA:*

ATGGACTACAAAGACGACGATGATAAGGCTAGCTCCTCCGACTCTCCGCGTAACCTGGAGGTTACCAACGCAACTCCG
AACTCTCTGACTATTTCTTGGGACAATTCTAACTATGCTTCGTATTACCGTATCACCTACGGCGAAACCGGTGGTAACT
CCCCGAGCCAGGAACTCACTGTTCCGGGAAGTACTTATAATGCGACCATCAGCGGTCTGAAACCGGGCCAGGATTATA
TCATTACCGTGTACGCTGTAACCTATCGTGACAATTATTCCTATTCAAATCTAATCAGCATCAATTATCGCTCCGAAATC
GACAAACCGTCTCAGGGATCCGGAGGTTCCGGCGGGGGCGGAAGCGGAGGTGGAGGCTCAGGGGGCGGAGGGTCGG
GCGGTGGAGGTTCGGGGGGAGGCGGGAGCGGTGGCGGTGGTTCAGGAGGAGGGGGTTCCGGGGGTGGTGGATCGGGC
GGTGAGCTCGGCGGCATCAGTCTGATTGCGGCGTTAGCGGTAGATCGCGTTATCGGCATGGAAAACGCCATGCCGTGG
AACCTGCCTGCCGATCTCGCCTGGTTTAAACGCAACACCTTAAATAAACCCGTGATTATGGGCCGCCATACCTGGGAAT
CAATCGGTCGTCCGTTGCCAGGACGCAAAAATATTATCCTCAGCAGTCAACCGGGTACGGACGATCGCGTAACGTGGG
TGAAGTCGGTGGATGAAGCCATCGCGGCGGCTGGTGACGTACCAGAAATCATGGTGATTGGCGGCGGTCGCGTTTATG
AACAGTTCTTGCCAAAAGCGCAAAAACTGTATCTGACGCATATCGACGCAGAAGTGGAAGGCGACACCCATTTCCCGG
ATTACGAGCCGGATGACTGGGAATCGGTATTCAGTGAATTCCACGATGCTGATGCGCAGAACTCTCACAGCTATAGCT
TTGAGATTCTGGAGCGGCGGGGCGGCATTAGCCTTATTGCCGCCTTAGCGGTTGATCGCGTGATCGGAATGGAGAACG
CAATGCCCTGGAATCTTCCGGCAGACCTTGCCTGGTTCAAACGCAACACTTTAAACAAGCCTGTCATTATGGGCCGTCA
CACATGGGAGTCAATTGGTCGTCCCCTGCCTGGGCGCAAAAATATCATCTTGTCCTCGCAGCCTGGGACAGATGATCG
CGTTACATGGGTGAAGTCCGTAGACGAAGCGATTGCCGCTGCCGGCGATGTGCCCGAGATTATGGTAATCGGGGGAGG
GCGTGTTTACGAACAATTTCTGCCCAAAGCTCAGAAATTATACCTGACGCACATCGACGCGGAGGTCGAAGGTGACAC
ACACTTTCCAGATTATGAGCCTGATGATTGGGAATCCGTTTTCTCAGAATTTCATGACGCGGATGCTCAAAACTCGCAC

TCGTACTCTTTTGAAATTTTAGAGCGCCGTGGCGGTTCAGGTGGTGGCTCGGGAGGCCATCATCATCATCATCACGGCG
GATCTGGAGGAAGTGGCGGTAAAAAGAAAAAGAAAAAGACCTGTGTCATCGCCTAGTGA

##### ***αCD133-scFv-DHFR²:***

ATGGACATTGTTCTCTCCCAGTCTCCAGCAATCATGTCTGCATCTCCAGGGGAGAAGGTCACCATATCCTGCAGTGCCA
GCTCAAGTGTAAGTTATATGTACTGGTACCAGCAGAAGCCAGGATCCTCCCCCAAACCCTGGATTTATCGCACATCCA
ACCTGGCTTCTGGAGTCCCTGCTCGCTTCAGTGGCAGTGGGTCTGGGACCTCTTACTCTCTCACAATCAGCAGCATGGA
GGCTGAAGATGCTGCCACTTATTACTGCCAGCAGTATCATAGTTACCCACCCACGTTCGGTGCTGGGACCAAGCTGGA
GCTGAAATCCTCTGGTGGCGGTGGCTCGGGCGGTGGTGGGGGTGGTTCCTCTAGATCTTCCCTCGAGGTGAAGCTGGT
GGAGTCTGGACCTGAGCTGAAGAAGCCTGGAGAGACAGTCAAGATCTCCTGCAAGGCTTCTGGTTATACCTTCACAGA
CTATTCAATGCACTGGGTGAATCAGGCTCCAGGAAAGGGTTTAAAGTGGATGGGCTGGATAAACACTGAGACTGGTGA
GCCATCATATGCAGATGACTTCAAGGGACGGTTTGCCTTCTCTTTGGAAACCTCTGCCAGCACTGCCTATTTGCAGATC
AACAACCTCAAAAATGAGGACACGGCTACATATTTCTGTGCTACCGATTACGGGGACTACTTTGACTACTGGGGCCAA
GGCACCACTCTCACAGTCTCCTCAGAGCTCGGCGGCAGCGGCGGCGGCAGCGGCGGCGGCAGCGGCGGCATGATCAG
TCTGATTGCGGCGTTAGCGGTAGATCGCGTTATCGGCATGGAAAACGCCATGCCGTGGAACCTGCCTGCCGATCTCGC
CTGGTTTAAACGCAACACCTTAAATAAACCCGTGATTATGGGCCGCCATACCTGGGAATCAATCGGTCGTCCGTTGCCA
GGACGCAAAAATATTATCCTCAGCAGTCAACCGGGTACGGACGATCGCGTAACGTGGGTGAAGTCGGTGGATGAAGC
CATCGCGGCGGCTGGTGACGTACCAGAAATCATGGTGATTGGCGGCGGTCGCGTTTATGAACAGTTCTTGCCAAAAGC
GCAAAAACTGTATCTGACGCATATCGACGCAGAAGTGGAAGGCGACACCCATTTCCCGGATTACGAGCCGGATGACTG
GGAATCGGTATTCAGTGAATTCCACGATGCTGATGCGCAGAACTCTCACAGCTATAGCTTTGAGATTCTGGAGCGGCG
GGGCATGATCAGTCTGATTGCGGCGCTAGCGGTAGATCGCGTTATCGGCATGGAAAACGCCATGCCGTGGAACCTGCC
TGCCGATCTCGCCTGGTTTAAACGCAACACCTTAAATAAACCCGTGATTATGGGGCGCCATACCTGGGAATCAATCGG
TAGGCCTTTGCCCGGCCGCAAAAATATTATCCTCAGCAGTCAACCCGGGACCGATGATCGGGTTACCTGGGTTAAATC
GGTCGACGAAGCCATCGCGGCGGCTGGTGACGTACCAGAAATCATGGTGATTGGCGGCGGACGCGTTTATGAACAGTT
CTTGCCAAAAGCGCAAAAACTGTATCTGACGCATATCGATGCAGAAGTGGAAGGCGACACCCATTTTCCGGATTACGA
GCCGGATGACTGGGAATCGGTATTCAGCGAGTTTCATGACGCGGACGCTCAGAACTCGCATAGCTATAGCTTCGAAAT
CCTGGAACGTCGTGGCGGCAGCGGCGGCGGCAGCGGCGGCGGCAGCGGCGGCGACTACAAAGACGATGACGACAAGT
AGTGA

##### ***DBCO-ssDNA-AF488:***

/5DBCOTEG/TGTCATATTCCTGGATCC/3AlexF488N/

##### ***DBCO-CTRLssDNA:***

TGTCATATTCCTGGATCCTTT/3DBCON/

#### ***DBCO-KDssDNA:***

T*G*T*C*A*T*A*T*T*C*C*T*G*G*A*T*C*C*T*T*T*/3DBCON/

(Note: * represents phosphorothioate bonds modifications of the backbone.)
